# Loss of FRIENDLY MITOCHONDRIA (FMT) Restores Chloroplast Proteostasis via Inter-organelle Compensation

**DOI:** 10.64898/2026.02.14.705898

**Authors:** Jitae Kim, Changhee Na, Pratyush Routray, NoA Bae, Hanseul Kim, Jae-Yeon Kim, Da Been Kim, Namil Son, Zeeshan Nasim, Ryoung Lee, Ji Hyun Kang, Geoni Choi, Hyoungseok Lee, Ji Hoon Ahn, Byeong-ha Lee, Dong Wook Lee, Klaas J. van Wijk

**Affiliations:** Department of Life Science, Sogang University, Seoul, 04107, Republic of Korea; Department of Integrative Food, Bioscience and Biotechnology, Chonnam National University, Gwangju, 61186, Republic of Korea; Department of Plant Biology, Cornell University, Ithaca, New York 14853, USA; Department of Plant Sciences, University of Cambridge, Cambridge CB2 3EA, UK; Department of Life Sciences, Korea University, Seoul 02841, Republic of Korea; Department of Applied Plant Science, Chonnam National University, Gwangju, 61186, Republic of Korea; Division of Life Sciences, Korea Polar Research Institute, Incheon, 21990, Republic of Korea; Department of Bioenergy Science and Technology, Chonnam National University, Gwangju, 61186, Republic of Korea

**Author notes:** Authors for correspondence (J.K.), (D.W.L.), (K.J.v.W.).

## Abstract

Chloroplast proteostasis is vital for plant development, yet whether cells can actively reprogram organelle communication to restore plastid function when essential protease components fail remains unclear. Using a forward genetic suppressor screen in *Arabidopsis*, we identify loss of the mitochondria-associated protein FRIENDLY MITOCHONDRIA (FMT) as a strong suppressor of the virescent and growth-retarded *clpc1* mutant, which lacks the major chloroplast Clp chaperone ClpC1. Suppression is highly specific and occurs independently of GUN1-mediated retrograde signaling. Integrated multi-omics analyses reveal that *clpc1* and *fmtclpc1* represent two distinct organelle signaling states. In *clpc1*, loss of ClpC1 triggers a plastid stress state characterized by repression of photosynthesis-associated transcription factors, induction of plastid metabolic stress markers, and impaired proteolytic activity. By contrast, loss of FMT shifts the system into a recovery state despite persistent mitochondrial clustering. Mechanistically, FMT negatively regulates *CLPC2*, a ClpC1 paralog, and *fmt*-mediated rescue results from *CLPC2* derepression. Moderately elevated ClpC2 restores *in vivo* proteolysis, as evidenced by recovery of PAA2 substrate turnover, normalization of chloroplast ultrastructure, and reactivation of photosynthesis-related gene expression. Transcriptomic and proteomic profiling further reveal coordinated remodeling of nuclear gene expression and chloroplast protein investment in the recovery state, including reduced cytosolic folding stress and selective induction of jasmonic acid– and salicylic acid–associated signaling networks. Genetic analyses establish that REC1 and REC2 are required for full *CLPC2* induction and phenotypic recovery. Together, our findings uncover a latent inter-organelle compensatory mechanism in which mitochondrial perturbation reprograms nuclear gene expression to restore chloroplast proteostasis when ClpC1 function is compromised.

## INTRODUCTION

Plant survival depends on the ability of its energy-producing organelles to communicate and maintain proteostasis despite continual developmental, metabolic, and environmental fluctuations (Gao et al., 2023; Sun and Jarvis, 2023). Chloroplasts and mitochondria rely on thousands of nuclear-encoded proteins that must be precisely synthesized, imported, and folded, rendering both organelles acutely vulnerable to even modest disruptions in protein homeostasis (Sun et al., 2021; Woodson and Chory, 2008). Because failures in these processes compromise photosynthesis, metabolism, and growth, plants must deploy coordinated cellular responses to proteotoxic stress across organelles (Llamas and Pulido, 2022; Lopez et al., 2024; van Wijk, 2024). Although plastid-to-nucleus retrograde signaling pathways that report organelle dysfunction have been extensively characterized, whether cells can transition from a stress-associated transcriptional state to an active recovery configuration that restores chloroplast proteostasis remains largely unknown.

In chloroplasts, proteostasis relies heavily on the stromal Clp chaperone–protease system, the most abundant and broadly acting proteolytic machinery in the organelle (Bouchnak and van Wijk, 2021; Rodriguez-Concepcion et al., 2019). The ClpPR core comprises catalytically active ClpP1 and ClpP3–6 and non-enzymatic ClpR1–4 subunits arranged as two stacked heptameric rings (Kim et al., 2013; Kim et al., 2009; Olinares et al., 2011; Sjogren et al., 2006). Substrate unfolding and translocation are mediated by the AAA+ chaperones ClpC1, ClpC2, and ClpD, while accessory proteins such as ClpT1/2, ClpS1, and ClpF contribute to core assembly/stabilization and substrate selection (Kim et al., 2015; Nishimura et al., 2015; Nishimura et al., 2013; Sjogren and Clarke, 2011). The turnover of numerous stromal proteins—including the canonical substrate PAA2/HMA8, a copper transporter in the thylakoid—highlights the central role of the Clp machinery in chloroplast protein quality control (Tapken et al., 2015).

ClpC1 is a pivotal component of this system. It functions both as the major unfoldase that delivers substrates to the ClpPR core (Rei Liao et al., 2022) and is also involved in quality control of imported proteins at the inner envelope membrane (Paila et al., 2015). Loss of ClpC1 leads to severe chloroplast dysfunction, manifesting as chlorosis, impaired photosynthesis, and markedly reduced growth. Protein-import stress is a known trigger of retrograde transcriptional reprogramming (Richter et al., 2023), and recent work has shown that GUN1 directly binds and stabilizes a subset of plastid RNAs, identifying plastid RNA maturation as a key upstream determinant of plastid-to-nucleus signaling (Tang et al., 2024). These studies define components of the plastid stress response; however, it remains unclear whether plants possess intrinsic mechanisms that can actively reconfigure nuclear gene expression to re-establish chloroplast proteostasis when a core chaperone such as ClpC1 is absent.

Mitochondria also undergo dynamic structural and functional remodeling in response to stress (Chustecki et al., 2022; Kacprzak and Van Aken, 2023; Nakamura et al., 2021). The conserved CLU-family protein FRIENDLY MITOCHONDRIA (FMT) maintains mitochondrial distribution, dynamics, and homeostasis; *fmt* mutants exhibit clustered mitochondria, altered organelle contacts, elevated ROS, and growth defects (El Zawily et al., 2014; Logan et al., 2003). FMT associates with cytosolic ribosomes at the mitochondrial surface to promote translation of nuclear-encoded mitochondrial mRNAs (Ayabe et al., 2021; Hemono et al., 2022), and contributes to mitophagy during de-etiolation (Ma et al., 2021). Despite these well-characterized mitochondrial functions, whether mitochondrial perturbation can influence plastid proteostasis or alter nuclear transcriptional states linked to chloroplast function has not been explored. Arabidopsis encodes three FMT-related homologues—*REC1-REC3*—that regulate chloroplast compartment size, and in tomato they additionally influence plastid development and plastid-related gene expression (Hu et al., 2024; Larkin et al., 2016), although their functions in organelle stress responses remain unresolved.

To identify pathways capable of actively restoring chloroplast proteostasis when ClpC1 function is lost, we performed a forward EMS suppressor screen in the severe chlorotic *clpc1-1* mutant. Multiple independent suppressors carried mutations in *FMT*, revealing an unexpected connection between mitochondrial perturbation and recovery of chloroplast function. RNA-seq and comparative proteomics uncovered coordinated remodeling of nuclear gene expression and chloroplast protein investment, consistent with a transition from a plastid stress state to a recovery state. We show that loss of *FMT* is associated with increased *REC2* expression and transcriptional upregulation of *CLPC2,* a ClpC1 paralog, and that *REC1* and *REC2* are required for full *fmt*-mediated compensation. Elevated ClpC2 functionally substitutes for ClpC1, restoring chloroplast proteostasis, pigmentation, and growth. Together, these findings reveal an inter-organelle compensatory mechanism in which mitochondrial perturbation enables reprogramming of nuclear gene expression to re-establish chloroplast proteostasis when ClpC1 function is compromised.

## RESULTS

### Suppressor Screen Identifies Green Leaf Mutants in the *clpc1-1* Background

To identify new genetic factors involved in chloroplast proteostasis and to gain insight into cellular responses to *CLPC1* deficiency, we performed a forward genetic suppressor screen in the *clpc1-1* background. The *clpc1-1* mutant exhibits a pronounced virescent and growth-retarded phenotype, enabling efficient visual identification of M2 individuals with restored greening and enhanced growth. Approximately 10,000 *clpc1-1* seeds were mutagenized with ethyl methanesulfonate (EMS), and the resulting M2 progeny were screened for recovery of the chlorotic phenotype. We identified ∼10 independent suppressor candidates showing partial or near-complete restoration of the wild-type phenotype. Among these, four lines with the strongest recovery—approaching wild type—were selected for detailed characterization (Figure 1A, B). The remaining candidates displayed weaker phenotypes and were not pursued further in this study.

**Figure 1.**
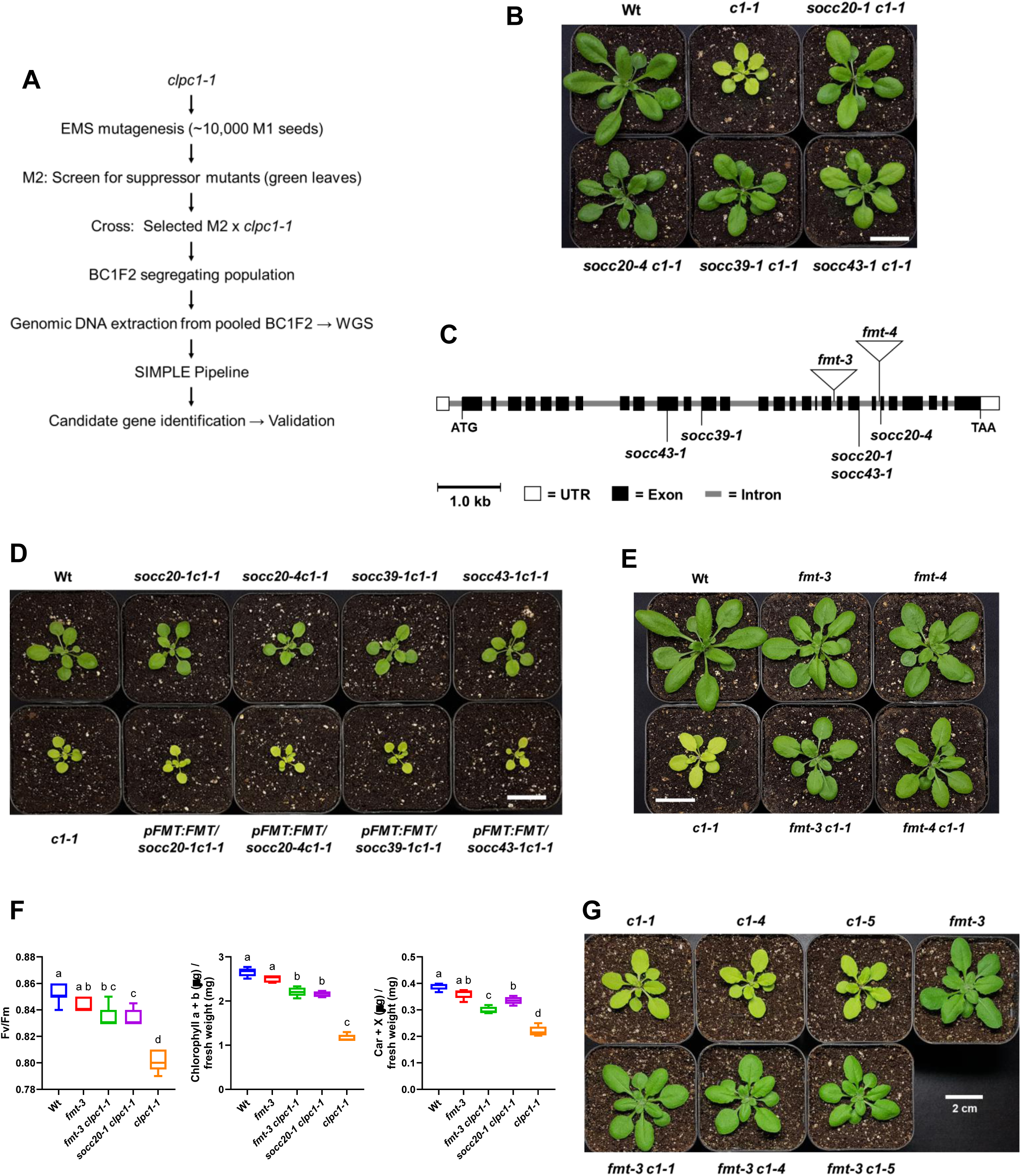
Identification and confirmation of *FMT* as the causal gene for suppressor mutations of *clpc1-1*. **(A)** Workflow of EMS mutagenesis, suppressor screening, and bulked segregant analysis. Seeds of *clpc1-1* were mutagenized with ethyl methanesulfonate (EMS) (∼10,000 seeds), and M2 plants were screened for suppressor mutants exhibiting a green-leaf phenotype. Selected M2 plants were backcrossed to the parental *clpc1-1* line (M0) to generate BC1F2 progeny. Pooled DNA from green BC1F2 plants was subjected to whole-genome sequencing (WGS), and mutations were identified using the SIMPLE pipeline (Wachsman et al., 2017). **(B)** Growth and development of wild-type (Col-0), *clpc1-1*, and four independent *clpc1-1* suppressor mutants - designated *socc* (<u>s</u>uppressor <u>o</u>f *<u>c</u>lp<u>c</u>1*) - including *socc20-1 clpc1-1*, *socc20-4 clpc1-1*, *socc39-1 clpc1-1*, and *socc43-1 clpc1-1*, grown on soil for 23 days under a 16/8 hour light/dark cycle at 100 µmol photons m⁻² s⁻¹. Scale bar = 2 cm. **(C)**Gene model of *FMT* (At3g52140) showing the positions of T-DNA insertions (*fmt-3*, *fmt-4*) and EMS-induced mutations identified in *socc* suppressor lines. Coding exons (black boxes), untranslated regions (open boxes), and introns (gray lines) are indicated. Mutation sites are mapped relative to the *FMT* genomic structure. The EMS-induced *socc* alleles include canonical splice-site defects and splice-region variants that disrupt *FMT* coding or splicing, while the T-DNA alleles (*fmt-3* and *fmt-4*) interrupt the gene’s exon–intron structure. These mutations affect *FMT* coding or splicing and are associated with the corresponding mutant phenotypes. Detailed mutation positions, nucleotide changes, and predicted effects are provided in table S1. **(D)** Genetic complementation of four suppressor mutants (*socc clpc1-1* lines) using a genomic *FMT* transgene driven by its native promoter. Transformation of each *socc clpc1-1* mutant with *pFMT:FMT* restored the yellow, chlorotic phenotype characteristic of *clpc1-1*, confirming *FMT* as the causal gene. Plants were grown on soil for 18 days under a 16/8 hour light/dark cycle at 100 µmol photons m⁻² s⁻¹. Scale bar = 2 cm. **(E)** T-DNA insertion alleles *fmt-3* and *fmt-4* suppress the *clpc1-1* phenotype, similar to EMS-derived suppressor lines. Growth and development of wild-type, *fmt-3*, *fmt-4*, *clpc1-1*, *fmt-3 clpc1-1*, and *fmt-4 clpc1-1* plants grown on soil for 23 days under a 16/8 hour light/dark cycle at 100 µmol photons m⁻² s⁻¹. Scale bar = 2 cm. **(E)** Photosynthetic efficiency and pigment accumulation in wild type, *fmt-3, clpc1-1*, and suppressor lines. Maximum quantum efficiency of PSII (Fv/Fm), total chlorophyll content, and total carotenoid content were measured in Col-0 (wild type), *fmt-3*, *clpc1-1*, *fmt-3 clpc1-1*, and *socc20-1 clpc1-1*. Fv/Fm measurements were performed on 22–24-day-old soil-grown plants, while pigment quantifications were conducted on 21-day-old plants, all grown under a 16/8 hour light/dark cycle at 100 μmol photons m⁻² s⁻¹. Fv/Fm values (left), total chlorophyll (middle), and total carotenoid contents (right) are shown. Chlorophyll and carotenoid contents are expressed as micrograms per milligram fresh weight (μg/mg FW). Bars represent mean ± SD (n = 7 for Fv/Fm; n = 5 for chlorophyll and carotenoid contents). Different lowercase letters indicate groups that differ significantly, as determined by one-way ANOVA followed by Tukey’s range test (P < 0.05). **(G)** Suppression of CRISPR/Cas9-derived *clpc1* mutant alleles (*clpc1-4* and *clpc1-5*) by *fmt-3*. Growth and development of wild-type, *clpc1-1*, *clpc1-4*, *clpc1-5*, *fmt-3*, *fmt-3 clpc1-1*, *fmt-3 clpc1-4*, and *fmt-3 clpc1-5* plants grown on soil for 20 days under a 16/8 hour light/dark cycle at 100 µmol photons m⁻² s⁻¹. Scale bar = 2 cm.

To genetically map the causal suppressor mutations, each of the four selected lines—designated *socc* (<u>s</u>uppressor <u>o</u>f *<u>c</u>lp<u>c</u>1*)—was backcrossed to the parental *clpc1-1* mutant. In the segregating BC1F2 populations, the green-leaf phenotype appeared at an approximate 1:3 ratio (green:chlorotic), consistent with suppression conferred by a single recessive locus. For each *socc* line, genomic DNA was extracted from more than 50 BC1F2 individuals displaying either green or chlorotic phenotypes. DNA from plants of the same phenotype was pooled and subjected to whole-genome sequencing. Candidate causal variants were then identified by comparing allele frequency differences between the green and chlorotic bulks using the SIMPLE pipeline (Wachsman et al., 2017) (Figure 1A).

### Whole-Genome Sequencing Reveals Recurrent Mutations in *FMT*

Analysis of single nucleotide polymorphism (SNP) frequencies across bulked segregant pools revealed that all four independent suppressor lines carried distinct EMS-induced mutations in a common gene, *FMT* (*At3g52140*). Representative phenotypes of these lines are shown in Figure 1B, each exhibiting substantial restoration of greening relative to the *clpc1-1* parental mutant.

In *socc43-1 clpc1-1*, a missense mutation in exon 10 of *FMT* results in a Glu534Lys substitution. In contrast, *socc20-1*, *socc20-4*, and *socc39-1* carry mutations affecting canonical splice donor or splice acceptor sites, or adjacent splice regions, all predicted to disrupt proper splicing and impair *FMT* function. Notably, *socc43-1* also harbors a synonymous substitution near an exon–intron boundary that may affect splicing efficiency (Figure 1C; Supplemental Table 1). The independent recovery of distinct EMS-induced mutations in *FMT* across multiple suppressor lines strongly implicates *FMT* as the candidate locus underlying suppression of *clpc1-1*.

### Complementation Confirms *FMT* as the Causal Suppressor Gene

To test whether loss of *FMT* is responsible for suppression, each of the four EMS-derived suppressor lines was transformed with a wild-type *FMT* genomic construct driven by its native promoter (*pFMT::FMT*). In all cases, reintroduction of functional *FMT* restored the chlorotic and stunted phenotype of *clpc1-1*, thereby reversing suppression (Figure 1D). These complementation results provide strong functional evidence that disruption of *FMT* is directly responsible for the green, growth-recovered phenotype of the suppressor lines.

### T-DNA Insertion Mutants Validate *FMT* as the Causal Suppressor Gene

To further validate *FMT* as the gene responsible for suppression of the *clpc1-1* phenotype, we analyzed two independent T-DNA insertion alleles, *fmt-3* and *fmt-4*. When combined with *clpc1-1*, both double mutants (*fmt-3clpc1-1* and *fmt-4clpc1-1*) exhibited restored greening and growth, phenocopying the EMS-derived suppressor lines (Figure 1E).

To assess the effects of EMS- and T-DNA–induced mutations on *FMT* transcript accumulation, we performed semi-quantitative RT-PCR using four primer sets spanning different regions of the *FMT* coding sequence (1F1R–4F4R) (Supplemental Figure 1A, B). In *fmt-3*, *FMT* transcript levels were strongly reduced across all primer sets, consistent with a severe knockdown. In contrast, *fmt-4* showed loss of amplification at the C-terminal region (3F3R and 4F4R), consistent with a null allele. EMS-derived *socc* alleles exhibited aberrant splicing patterns, with *socc20-1*, *socc20-4*, and *socc43-1* producing additional higher-molecular-weight bands with the 3F3R primer set, indicative of mis-spliced transcripts. Across all EMS alleles tested, overall *FMT* transcript abundance was reduced relative to *clpc1-1* (Supplemental Figure 1B), consistent with partial loss of function.

As expected, *CLPC1* transcripts were absent in all *clpc1-1* backgrounds, confirming that *clpc1-1* is a null allele and that suppression does not arise from *CLPC1* reactivation. Notably, *FMT* transcripts were elevated in *clpc1-1* relative to wild type, suggesting transcriptional upregulation of *FMT* in response to chloroplast dysfunction caused by *CLPC1* loss. Together, these data validate the molecular identity of both T-DNA and EMS-derived suppressor alleles and demonstrate that suppression arises through loss of *FMT* function rather than restoration of *CLPC1*.

To test whether *fmt*-mediated suppression restores chloroplast performance, we measured maximum quantum efficiency of PSII (Fv/Fm) and total pigment content. Both *fmt-3clpc1-1* and *socc20-1clpc1-1* displayed significantly higher Fv/Fm values than *clpc1-1*, approaching wild-type levels (Figure 1F, left). Total chlorophyll and carotenoid contents showed a similar trend (Figure 1F, middle and right). These results indicate that *fmt*-mediated suppression improves chloroplast function—enhancing photosynthetic efficiency and pigment accumulation in the *clpc1-1* background.

### Heteroallelic Crosses Confirm *FMT* as the Causal Suppressor Gene

To confirm that the EMS-derived *socc* suppressor mutations disrupt *FMT* rather than other loci, we performed heteroallelic complementation tests. F1 progeny from *fmt-3clpc1-1×fmt-4clpc1-1* crosses exhibited the same suppressed phenotype as the parental lines, confirming allelism between the two T-DNA insertions (Supplemental Figure 2A). Likewise, F1 progeny from each EMS-derived *socc_clpc1-1×fmt-3clpc1-1* cross displayed the suppressed phenotype, indicating failure to complement. These results establish that all four *socc* alleles represent independent loss-of-function mutations in *FMT*.

### CRISPR-Derived *clpc1* Null Alleles Confirm Allele-Independent Suppression by *fmt*

To rule out T-DNA–specific epigenetic suppression of *clpc1-1*, we generated two independent CRISPR-derived null alleles (*clpc1-4*, *clpc1-5*), each carrying frameshift mutations in exon 4. Both alleles phenocopied *clpc1-1* and failed to complement it in heteroallelic crosses (Supplemental Figure 2B). Importantly, both were robustly suppressed by *fmt-3* (Figure 1G), demonstrating that *fmt*-mediated rescue is allele-independent and does not result from epigenetic reactivation (Gao and Zhao, 2013; Jia et al., 2015) of the T-DNA insertion. Together, these results establish *FMT* loss of function as the causal basis of all recovered EMS suppressors and demonstrate that disruption of *FMT* restores chloroplast performance in multiple *clpc1* null alleles through a genuine genetic interaction with *CLPC1*.

### *fmt* Specifically Suppresses *clpc1-1* but Not Other Chloroplast Proteostasis or Import Mutants

We next investigated whether *fmt*-mediated suppression is specific to *CLPC1* or extends to other components of chloroplast proteostasis and protein import. To test this, we examined genetic interactions between *fmt* and representative mutants affecting distinct subunits of the Clp protease complex and the chloroplast import machinery.

To determine whether suppression extended to other components of the Clp chaperone-protease complex, we crossed *fmt* with *clpr2-1* (Rudella et al., 2006), a non-catalytic core subunit mutant, and with the *clpt1-2clpt2-1* double mutant (Kim *et al*., 2015), which lacks accessory proteins critical for Clp complex stability and assembly. Neither *fmt-3* nor *fmt-4* suppressed the pale, small phenotype of *clpr2-1* (Figure 2A), and *fmt-4* did not rescue the severe developmental defects of *clpt1-2clpt2-1* (Figure 2B). These results indicate that *fmt* suppression is not broadly applicable to defects in the Clp protease complex and may act specifically at the level of ClpC1.

**Figure 2.**
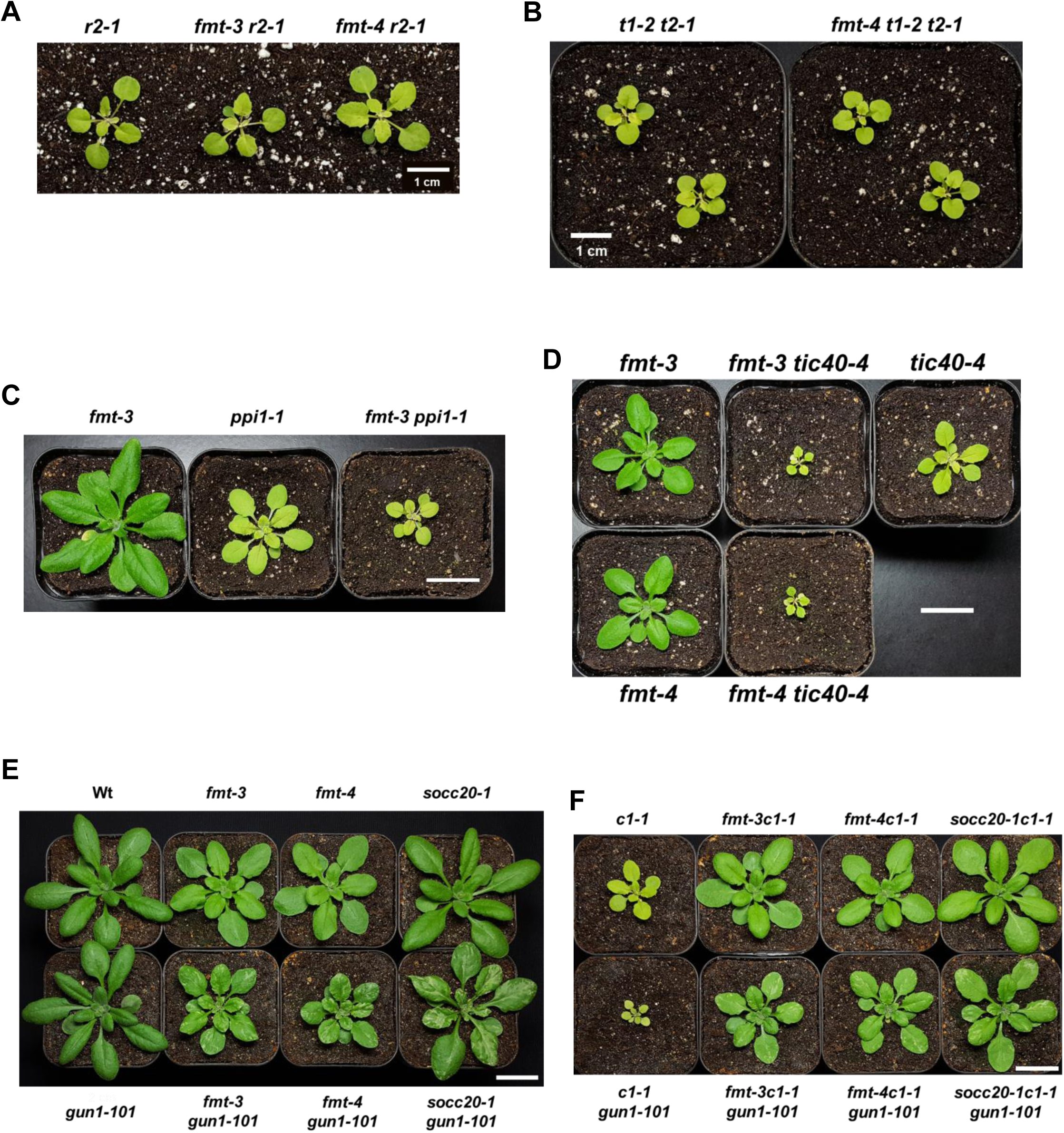
Genetic Interaction of various mutants with *fmt*. **(A)** Growth and development of *clpr2-1, fmt-3 clpr2-1,* and *fmt-4 clpr2-1* mutants. Plants were grown on soil for 22 days under a 16/8 hour light/dark cycle at 100 µmol photons m⁻² s⁻¹. Scale bar = 1 cm. **(B)** Growth and development of *clpt1-2 clpt2-1* and *fmt-4 clpt1-2 clpt2-1* mutants. Plants were grown on soil for 29 days under the same conditions. Scale bar = 1 cm. **(C)** Growth and development of *fmt-3, ppi1-1,* and *fmt-3 ppi1-1* mutants. Plants were grown on soil for 24 days under the same conditions. Scale bar = 2 cm. **(D)** Growth and development of *fmt-3, fmt-4, tic40-4, fmt-3 tic40-4,* and *fmt-4 tic40-4* mutants. Plants were grown on soil for 20 days under the same conditions. Scale bar = 2 cm. **(E)**Growth and development of wild-type (Col-0), *fmt-3*, *fmt-4*, *socc20-1*, *gun1-101*, *fmt-3 gun1-101*, *fmt-4 gun1-101*, and *socc20-1 gun1-101* plants grown on soil for 21 days under a 16/8 hour light/dark cycle at 100 µmol photons m⁻² s⁻¹. Variegated phenotypes were observed in all *fmt gun1-101* double mutants. Scale bar = 2 cm. **(F)** Growth and development of *clpc1-1*, *fmt-3 clpc1-1*, *fmt-4 clpc1-1*, *socc20-1 clpc1-1*, *clpc1-1 gun1-101*, *fmt-3 clpc1-1 gun1-101*, *fmt-4 clpc1-1 gun1-101*, and *socc20-1 clpc1-1 gun1-101* plants grown on soil for 21 days under a 16/8 hour light/dark cycle at 100 µmol photons m⁻² s⁻¹. Suppression of the *clpc1-1 gun1-101* growth defect was observed in all *fmt clpc1-1 gun1-101* triple mutants. Scale bar = 2 cm.

We next tested genetic interactions between *fmt* and mutants defective in the chloroplast protein import. The *ppi1-1* mutant, lacking the TOC33 receptor and displaying a pale-green, small size seedling phenotype (Kubis et al., 2003), was crossed with *fmt-3*. Similarly, the small and pale *tic40-4* mutant, defective in a co-chaperone of the TIC complex (Bedard et al., 2017), was crossed with both *fmt-3* and *fmt-4*. In all cases, the resulting double mutants displayed exacerbated phenotypes relative to the respective single mutants. Specifically, *fmt-3ppi1-1* plants were smaller than *ppi1-1* alone, and *fmt-3tic40-4* and *fmt-4tic40-4* plants were markedly smaller and more chlorotic than *tic40-4* (Figure 2C, D). These results show that *fmt* does not restore defects in chloroplast protein import and that loss of *FMT* further impairs plastid function when import is compromised. Thus, *fmt*-mediated suppression is highly specific to *clpc1-1* and does not reflect general rescue of Clp or import defects.

### *fmt*-mediated suppression of *clpc1-1* operates independently of GUN1

To examine whether *fmt* interacts with plastid-to-nucleus retrograde signaling pathways (Richter *et al*., 2023), we combined *fmt* alleles with *gun1-101*. While *fmt* and *gun1-101* single mutants were uniformly green, all *fmtgun1-101* double mutants displayed a distinctive variegated phenotype (Fig. 2E), indicating that loss of *GUN1* results in unstable chloroplast biogenesis and function in *fmt* backgrounds.

We next tested whether *GUN1* is required for *fmt*-mediated suppression of *clpc1-1*. The *clpc1-1gun1-101* double mutant exhibited a more severe phenotype than *clpc1-1* alone (Figure 2F). However, introduction of *fmt-3*, *fmt-4*, or *socc20-1* into the *clpc1-1gun1-101* background restored greening and growth (Figure 2F), demonstrating that suppression occurs in the absence of *GUN1*. Notably, the triple mutants retained the variegation characteristic of *fmt gun1-101*, suggesting that loss of ClpC1 and loss of GUN1 have different underlying molecular causes (Figure 2F).

### *fmt* Suppressor Lines Restore Chloroplast Ultrastructure in *clpc1-1* Despite Persistent Mitochondrial Clustering

Previous studies showed that wild-type chloroplasts exhibit compact morphology with well-stacked thylakoids, whereas *fmt* mutants retain normal chloroplast ultrastructure but display abnormal mitochondrial clustering and aggregation (El Zawily *et al*., 2014). Consistent with its severe chlorotic phenotype, *clpc1-1* chloroplasts appeared swollen and irregular, with reduced stromal density and poorly organized thylakoid membranes (Figure 3A). In contrast, *fmt-3clpc1-1* double mutants exhibited compact chloroplasts with dense stroma and well-developed grana stacks, closely resembling wild type (Figure 3A). Mitochondria in the double mutants remained tightly clustered, reflecting the characteristic *fmt* phenotype.

**Figure 3.**
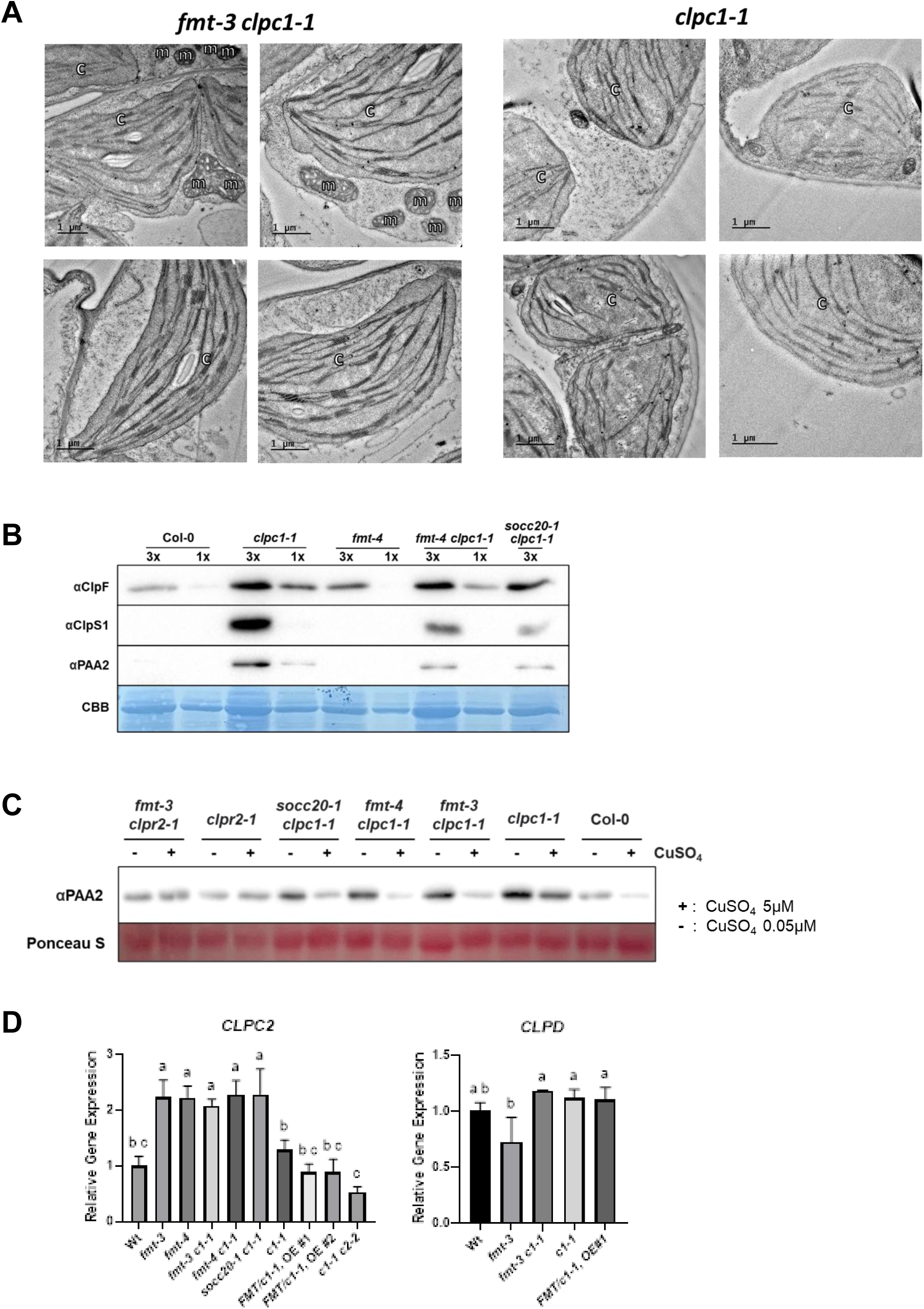
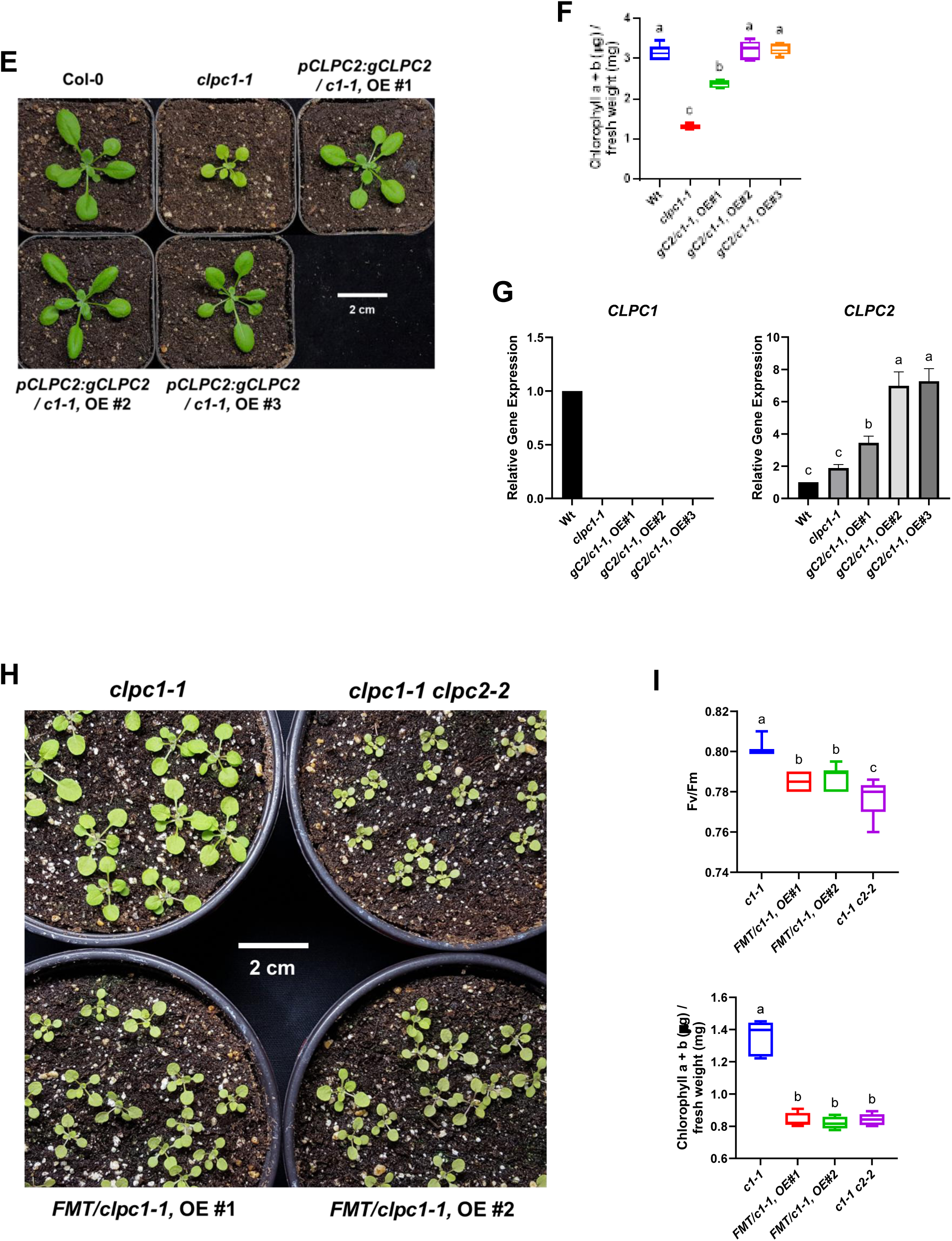
Loss of FMT restores chloroplast morphology and protease activity through compensatory upregulation of *CLPC2*. **(A)** Chloroplast ultrastructure in *fmt-3clpc1-1* and *clpc1-1*. Left panel: Transmission electron microscopy (TEM) of chloroplasts from *fmt-3 clpc1-1* leaf mesophyll cells. Chloroplasts exhibit compact morphology, well-organized grana stacks, and dense stromal regions, closely resembling wild type. Notably, mitochondria remain tightly clustered, consistent with the known *fmt* phenotype. Scale bar = 1 µm. Right panel: TEM of chloroplasts from *clpc1-1*. Chloroplasts appear swollen and rounded, with fewer and less organized thylakoid membranes and smaller, loosely stacked grana. Stromal regions are less compact than in the double mutant. Scale bar = 1 µm. m, mitochondria; c, chloroplast. **(B)** Immunoblot analysis of selected chloroplast proteins in Col-0, *clpc1-1*, *fmt-4*, *fmt-4 clpc1-1*, and *socc20-1 clpc1-1*. Antibodies used: ClpF (a Clp N-recognin adaptor protein), ClpS1 (a substrate selector for the Clp complex), and PAA2 (a Clp substrate involved in thylakoid copper transport). Each lane was loaded with either 10 µg (1×) or 30 µg (3×) of total chloroplast protein. Coomassie Brilliant Blue (CBB) staining shows equal loading. **(C)**PAA2 degradation assay using total protein extracts treated with or without CuSO₄. In Col-0, Cu²⁺ exposure leads to decreased PAA2 levels, whereas in *clpc1-1*, PAA2 remains elevated regardless of treatment. Cu²⁺-dependent degradation is restored in *fmt-3 clpc1-1*, *fmt-4 clpc1-1*, and *socc20-1 clpc1-1*. No degradation is observed in *fmt-3 clpr2-1*. Ponceau S staining confirms equal loading. **(D)** Quantitative RT-PCR analysis of *CLPC2* transcript levels in wild type (Col-0), *fmt-3, fmt-4*, *fmt-3 clpc1-1, fmt-4 clpc1-1*, *socc20-1 clpc1-1*, *clpc1-1*, *FMT/clpc1-1* overexpression (OE) lines (#1–2), and *clpc1-1 clpc2-2*. For comparison, *CLPD* transcript levels are shown for wild type (Col-0), *fmt-3*, *fmt-3 clpc1-1*, *clpc1-1*, and *FMT/clpc1-1* OE#1. Data represent means ± SD from three biological replicates. **(E)** Growth phenotype of *clpc1-1* and independent *pCLPC2:gCLPC2/clpc1-1* overexpression (OE) lines (#1–3) grown on soil for 18 days under a 16/8 hour light/dark cycle at 90 µmol photons m⁻² s⁻¹. Scale bar = 2 cm. **(F)** Total chlorophyll (a + b) accumulation in the lines shown in (E). Chlorophyll contents were quantified in 25-day-old plants grown under the same conditions and are expressed as micrograms per milligram fresh weight (µg/mg FW). Values represent mean ± SD (n = 5). **(G)** RT–qPCR analysis of *CLPC1* and *CLPC2* transcript levels in the same lines shown in (E). OE lines with higher *CLPC2* expression (e.g., OE #2 and OE #3) exhibit a greener phenotype compared with OE #1. Bars represent mean ± SD (n = 3). **(H)** Growth and development of *clpc1-1*, *clpc1-1 clpc2-2*, and *FMT/clpc1-1* plants grown on soil for 17 days under a 16/8 hour light/dark cycle at 100 µmol photons m⁻² s⁻¹. Both *clpc1-1 clpc2-2* and *FMT/clpc1-1* plants exhibited severely stunted and chlorotic phenotypes compared to *clpc1-1*. Overexpression lines were slightly larger than *clpc1-1 clpc2-2*. **(I)** Photosynthetic efficiency and total chlorophyll (a + b) accumulation in the same lines shown in (H). Maximum quantum efficiency of PSII (Fv/Fm; upper) and total chlorophyll contents (lower) were measured in 24-day-old soil-grown plants grown under a 16/8 hour light/dark cycle at 100 μmol photons m⁻² s⁻¹. Chlorophyll contents are expressed as micrograms per milligram fresh weight (μg/mg FW). Bars represent mean ± SD (n = 15 for Fv/Fm; n = 5 for chlorophyll contents). (D), (F), (G) and (I) Different lowercase letters indicate groups that differ significantly, as determined by one-way ANOVA followed by Tukey’s range test (P < 0.05).

### *fmt* Suppressor Lines Normalize ClpC1-Dependent Substrate Selector and Substrate Levels

To assess whether *fmt*-mediated suppression restores chloroplast proteostasis, we examined steady-state levels of ClpC1-associated proteins by immunoblotting (Figure 3B). ClpF (an N-recognin adaptor), ClpS1 (a substrate selector), and PAA2 (a thylakoid Cu-transporting ATPase and Clp substrate) were analyzed as representative markers of ClpC1-dependent function. PAA2 degradation requires ClpC1 under copper-replete conditions (Tapken *et al*., 2015), making it a sensitive marker of ClpC1-dependent activity. Immunoblotting showed that *clpc1-1* accumulated high levels of ClpF, ClpS1, and the ClpC1 substrate PAA2 (Fig. 3B), consistent with previous reports (Nishimura *et al*., 2015; Nishimura *et al*., 2013; Tapken *et al*., 2015). In *fmt-4clpc1-1* and *socc20-1clpc1-1*, ClpS1 and PAA2 levels decreased toward wild-type abundance, and ClpF levels trended downward.

### *fmt* Suppressor Lines Restore Clp Protease Activity

To directly assess whether suppression restores functional proteolysis, we performed a PAA2 degradation assay under CuSO₄-replete conditions, following the approach of (Tapken *et al*., 2015). PAA2 (HMA8) is a thylakoid Cu-transporting ATPase that is selectively degraded under high Cu²⁺ conditions in a ClpC1-dependent manner. Arabidopsis seedlings were grown on MS agar plates with or without CuSO₄, and PAA2 abundance was analyzed from total protein extracts (Figure 3C). In wild-type, CuSO₄ treatment caused a pronounced decrease in PAA2 levels, consistent with active Clp-mediated degradation. In *clpc1-1*, PAA2 remained highly abundant and insensitive to CuSO₄, indicating a strong defect in proteolytic turnover. Strikingly, Cu²⁺-dependent PAA2 degradation was restored in all three *fmt*-mediated suppressor lines: *fmt-3clpc1-1*, *fmt-4clpc1-1*, and *socc20-1clpc1-1*. In each case, CuSO₄ treatment led to a marked reduction in PAA2 levels, closely resembling the wild-type response and demonstrating that *fmt* suppression restores Cu²⁺-dependent Clp protease activity. In contrast, *fmt-3clpr2-1* exhibited persistent PAA2 accumulation similar to *clpr2-1*, reinforcing that *fmt* selectively suppresses *clpc1-1* but does not bypass defects in the Clp protease core.

### *fmt*-Mediated Suppression of *clpc1-1* Involves Transcriptional Upregulation of *CLPC2*

The recovery of chloroplast structure and proteostasis in *fmtclpc1-1* mutants (Figure 3A, C) suggested that *fmt* may activate a compensatory mechanism that restores Clp chaperone capacity in the absence of ClpC1. Because ClpC1 and ClpC2 have partially overlapping functions—with ClpC2 able to substitute for ClpC1 when expressed at higher levels—we asked whether *fmt* suppressors increase *CLPC2* expression.

To test this, we quantified *CLPC2* transcript abundance, along with *CLPD* as a related control, by RT–qPCR across multiple genotypes (Figure 3D). *CLPC2* expression was approximately twofold higher in *fmt-3*, *fmt-4*, *fmt-3clpc1-1*, *fmt-4clpc1-1*, and *socc20-1clpc1-1* relative to wild type and *clpc1-1*, indicating that loss of *FMT* consistently induces *CLPC2* transcription. In contrast, *CLPD* transcript levels showed only minor variation among genotypes and did not exhibit the coordinated upregulation seen for *CLPC2*, indicating that the compensatory response is specific to *CLPC2* rather than a general induction of Clp chaperone genes.

### Native Promoter–Driven Elevation of *CLPC2* Is Sufficient to Rescue the *clpc1-1* Phenotype

Because *CLPC1* expression is normally far higher than *CLPC2* in wild-type plants, we next asked whether the moderate increase in *CLPC2* observed in *fmtclpc1-1* suppressor lines is sufficient to account for phenotypic rescue. To test this, and to avoid artifacts associated with strong constitutive promoters, we generated *pCLPC2::genomic-CLPC2/clpc1-1* lines expressing genomic *CLPC2* under its native promoter.

These lines showed 2-4-fold increases in *CLPC2* transcripts compared with *clpc1-1* and showed proportional restoration of greening and pigment accumulation (Fig. 3E–G) (*i.e.* higher *CLPC2* transcript levels result in better suppression/complementation), demonstrating that a several-fold increase in ClpC2 is enough to offset complete loss of *clpc1-1*. The scaling of *CLPC2* expression across the overexpression lines is consistent with differences in *gCLPC2* copy number, and indicates that the promoter is not saturated in the *clpc1-1* background.

Together, these findings indicate that physiologically modest upregulation of *CLPC2* is sufficient to compensate for loss of *CLPC1* function. This finding complements earlier work by (Kovacheva et al., 2007), who showed that strong *35S::CLPC2* overexpression can also rescue *clpc1* defects, although high expression occasionally triggered co-suppression-associated albino phenotypes.

### *FMT* Expression Level Negatively Correlates with Growth and Chlorosis Severity in *clpc1-1*

To further examine how *FMT* expression levels influence the *clpc1-1* phenotype, we generated *pFMT::FMT/clpc1-1* lines in which *FMT* was expressed at elevated levels under its native promoter, minimizing non-physiological expression patterns sometimes associated with the 35S promoter. Seven independent OE lines were obtained, spanning a broad range of rosette size and chlorosis severity (Supplemental Figure 3A). Notably, higher *FMT* expression consistently correlated with smaller rosettes and reduced greening. Lines with the strongest overexpression (e.g., OE#1) displayed the most pronounced chlorosis, whereas lines with lower expression (e.g., OE#7) more closely resembled *clpc1-1*. Semi-quantitative RT-PCR confirmed graded differences in *FMT* transcript abundance among these lines (Supplemental Figure 3B), and densitometric analysis revealed a clear inverse relationship between *FMT* transcript level and phenotype severity (Supplemental Figure 3C).

Based on this expression gradient, we selected OE#1 and OE#2—the two lines with the highest *FMT* transcript accumulation—for detailed physiological and molecular characterization (Figure 3H–I).

### Overexpression of *FMT* Represses *CLPC2* and Exacerbates the *clpc1-1* Phenotype

We noticed that the smaller, chlorotic rosette phenotype of *pFMT::FMT/clpc1-1* overexpression lines (OE#1 and OE#2) closely resembled that of the *clpc1-1clpc2-2* double mutant (Figure 3H). *clpc2-2* is a partial loss-of-function allele, and *clpc1-1clpc2-2* plants exhibit a smaller, more chlorotic phenotype than *clpc1-1*, whereas the *clpc2-3* null mutant causes embryo lethality in the *clpc1-1* background (Nishimura *et al*., 2013). Although the two *FMT* overexpression lines were slightly larger than *clpc1-1clpc2-2*, they were markedly smaller and paler than *clpc1-1*, suggesting that elevated *FMT* expression exacerbates the *clpc1-1* phenotype by repressing—rather than supporting—*CLPC2*.

To verify the molecular basis of this phenotype, we quantified *CLPC2* transcript levels by RT-qPCR (Figure 3D). *CLPC2* transcript abundance in the *pFMT::FMT/clpc1-1* overexpression lines was approximately 60-70% of *clpc1-1* levels, while the *clpc1-1clpc2-2* double mutant retained about 50% of *clpc1-1* expression (Figure 3D). The lower *CLPC2* expression in *clpc1-1clpc2-2* was consistent with its more severe phenotype (Figure 3H). These results demonstrate that *FMT* overexpression represses *CLPC2* transcription in the *clpc1-1* background, supporting the conclusion that *FMT* functions as a negative regulator of *CLPC2*.

To determine whether these transcriptional differences impact chloroplast performance, we measured the maximum quantum yield of PSII (Fv/Fm) and pigment content (Figure 3I). Both *FMT* OE lines and *clpc1-1clpc2-2* exhibited significantly reduced Fv/Fm and pigment levels relative to *clpc1-1*, consistent with their more chlorotic and stunted phenotypes.

Together, these findings indicate that *fmt*-mediated suppression of *clpc1-1* involves transcriptional upregulation of *CLPC2*, which compensates for loss of ClpC1. Conversely, elevated *FMT* expression reduces *CLPC2* transcript abundance and phenocopies partial *CLPC2* deficiency, exacerbating the *clpc1-1* phenotype.

### RNA-seq reveals coordinated transcriptional remodeling associated with *clpc1* suppression

To better understand the molecular phenotypes of *fmt* and *clpc1-1,* and suppression of the *clpc1-1* phenotype in *fmtclpc1-1*, we performed RNA-seq analysis of Arabidopsis seedlings at developmental stage 1.07 for WT, *fmt-3, clpc1-1*, *fmt-3clpc1-1* and *FMT/clpc1-1* (OE#1). After mapping, quality control, and normalization, mRNA expression values were obtained for 21,202 genes (Supplemental Data Set 1A), and pairwise differential expression statistics were calculated for downstream analyses. Principal Component Analysis (PCA) showed a very distinct signature for *clpc1,* which was further enhanced in *FMT*/*clpc1* (Fig. 4A). *fmtclpc1* was more closely related to WT, whereas *fmt* was very similar to WT but remained distinct from *fmtclpc1*. The PCA results are consistent with the observed phenotypes across these five genotypes.

**Figure 4.**
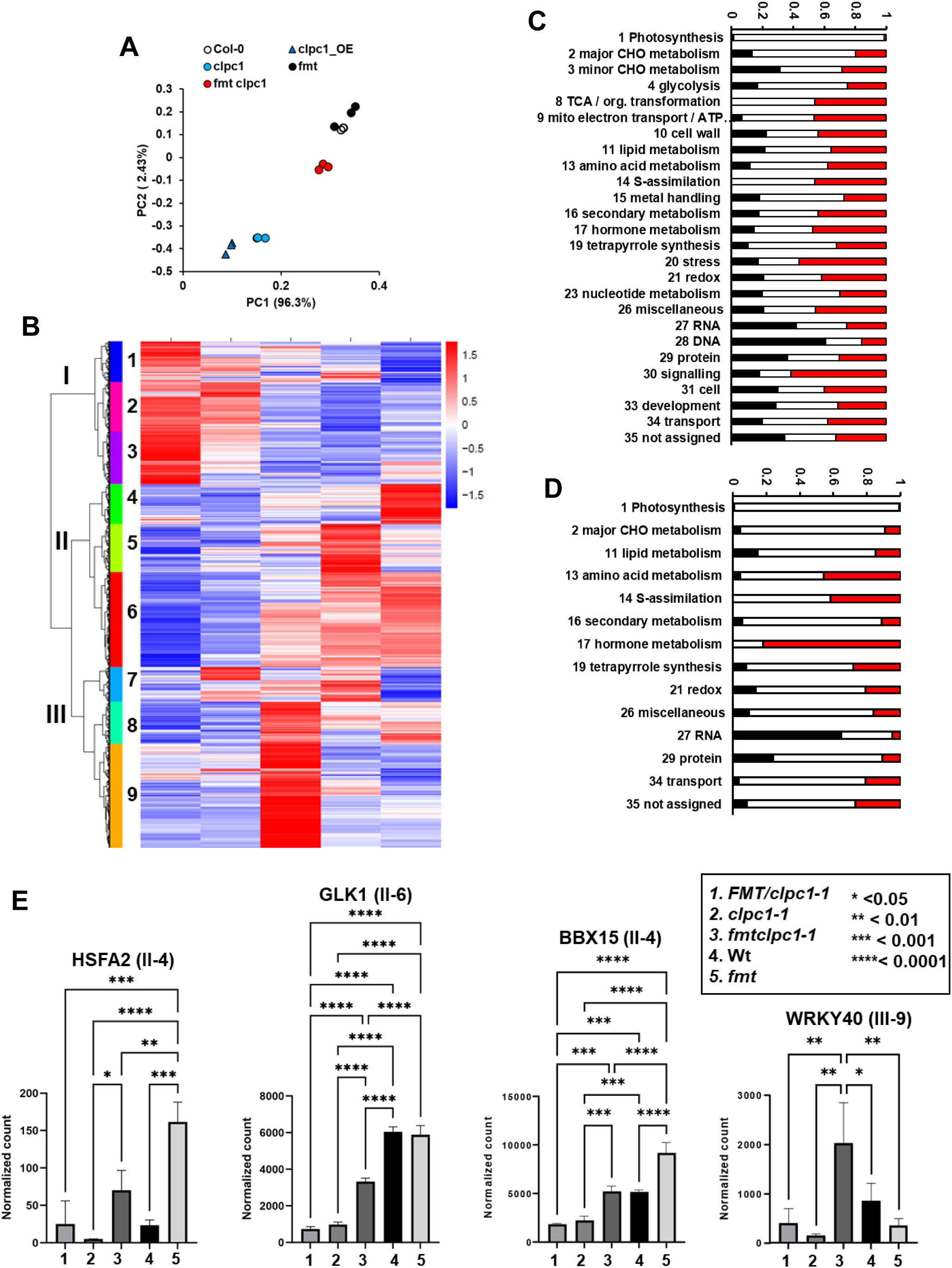
RNA expression patterns across the five genotypes wild type (Col-0), *fmt, clpc1, fmtclpc1* and *FMT/clpc1*. **(A)** Principal Component Analysis (PCA) of RNA-seq data showing clustering of biological replicates by genotype. **(B)** Hierarchical cluster analysis of z-score-normalized average RNA abundance of 4745 protein-coding nuclear genes in WT*, fmt, clpc1, fmtclpc1* and *FMT/clpc1* sampled at stage 1.07. Only genes that show at least one pairwise significance difference (adj P<0.01 and at least 2-fold difference) between 2 genotypes are included. The cladogram has three main clusters (I, II, III) with three subclusters each. The scaling of heat map is shown in the top right. The z-score was calculated as (individual value-average)/standard deviation for each gene across the 5 genotypes. **(C)** M apMan functional bin distribution across clusters I, II and III for all genes in the cladogram (4745 genes). Only bins with more than 10 genes are shown. **(D)** MapMan functional bin distribution of plastid and mitochondrial proteins across clusters I, II and III (638 genes). Only bins with more than 10 genes are shown. **(E)** Differential expression of the transcription factors **HSFA2, GLK1, BBX15, and WRKY40**. RNA-seq–normalized counts are shown as means ± SD for each genotype. Statistical significance was assessed using ordinary one-way ANOVA with Tukey’s multiple-comparison test. Significant differences are indicated by asterisks (\**P* < 0.05, \*\**P* < 0.01, \*\*\**P* < 0.001, \*\*\*\**P* < 0.0001). Gene cluster membership is indicated.

We extracted all genes that showed a pairwise difference of at least 2-fold (x<0.5 or x>2) with P<0.01 in at least one of the 10 possible comparisons across genotypes (Supplemental Data Set 1B). This resulted in 4,745 nuclear protein-coding genes that we annotated for subcellular protein location and function (Supplemental Data Set 1B). Gene cluster analysis showed three main clusters (I, II, III) that each further separated into three subclusters (Fig. 4B). Cluster I is dominated by genes for which the mRNA is enriched in *clpc1* and *FM*T/*clpc1* often with most enhanced expression in *FMT/clpc1*. Cluster II is enriched for genes that are reduced in *clpc1* and *FMT/clpc1.* Cluster III contains genes with mRNA generally enriched only in *fmtclpc1* as compared to the other four genotypes. MapMan (Thimm et al., 2004) functional bin analysis shows that cluster I and III are both highly depleted for genes encoding photosynthetic proteins (Fig. 4C). Cluster I, but not clusters II and III, is also relatively low in genes involved in mitochondrial electron transport/ATP synthesis (bin 9), tetrapyrrole metabolism (bin 19) and amino acid metabolism (bin 13). Cluster I is most enriched for genes involved in DNA and RNA related processes. Most genes in cluster III (subcluster 8 and 9) are expressed much higher in *fmtclpc1* than in any of the other four genotypes; this cluster is especially enriched for signaling (bin 30) and stress (bin 20) (Fig. 4C). Figure 4D shows the bin-cluster analysis for the genes encoding experimentally confirmed plastid-, mitochondrial-, and dually localized proteins (556, 58 and 15 genes, respectively). 72% belong to cluster II. RNA metabolism shows strong enrichment for cluster I – many of these are PPR proteins with functions in editing and RNA processing of plastid- or mitochondria-encoded genes. Proteins involved in the protein life cycle (bin 29) are also enriched in this cluster I. Photosynthetic functions (bin 1) and starch metabolism (bin 2) are nearly exclusively in cluster II, consistent with the loss of photosynthesis phenotypes of *clpc1* and *FMT/clpc1*. In contrast, amino acid metabolism (bin 13), sulfur assimilation (bin 14), and hormone synthesis (bin 17 - nearly exclusively proteins involved in plastid jasmonate metabolism, including allene oxide cyclase 1, 2, 3, and 4; allene oxide synthase; and lipoxygenase 1 and 2) are enriched in cluster III.

Subcluster 4 (376 genes) stands out for its high mRNA levels in the *fmt* line compared to the other four genotypes. Many of the genes with the most pronounced expression profiles encode mitochondrial proteins (Supplemental Figure 4A), including alternative oxidase 1A (AOX1A), stomatin-like protein 2 (SLP2), heat shock protein HSP23.5, DUF295 domain protein DOB5, and external NAD(P)H dehydrogenase subunit B3 (NDH-B3), all of which are reported to have low basal expression levels (Elhafez et al., 2006). AOX1A and NDH-B3 partially uncouple respiration from mitochondrial ATP production and thereby reduce mitochondrial reactive oxygen species (ROS) production (Selinski et al., 2018). AOX1A is a marker of the mitochondrial retrograde response (Wang et al., 2020). SLP2 affects protein abundance and activity of respiratory complex I, likely through effects on its assembly and/or stability (Gehl et al., 2014; Gehl and Sweetlove, 2014). HSP23.5 is a member of the HSP20-like chaperone superfamily; although this specific protein has not been characterized, mitochondrial HSP20 chaperones are important for facilitating protein folding within mitochondria, particularly under stress conditions (Waters and Vierling, 2020). DUF295 domain protein DOB5 has not been characterized but is part of the large DUF295 family (Lama et al., 2019). Notably, subcluster 4 contains eight DUF295 family members, seven of which are predicted to encode mitochondrial proteins. Many mitochondrial DUF295 proteins are induced by mitochondrial dysfunction (Lama *et al*., 2019), which is highly consistent with their elevated abundance in the *fmt* background.

Subclusters eight and nine (within cluster III) contain genes with high mRNA levels in *fmtclpc1* compared to the other four genotypes. These genes may provide clues about the nature of *clpc1* suppression by *fmt.* Among the confirmed plastid and mitochondrial proteins, we identified five genes for which the RNA levels in *fmtclpc1* were more than threefold higher than in each of the other four genotypes (Supplemental Figure 4B). Surprisingly, all five are involved in jasmonate or salicylic acid metabolism or response; these include two allene oxide cyclases (AOC1 and AOC3) and lipoxygenase 3 involved in JA synthesis (Bittner et al., 2022), the plastid envelope MATE transporter EDS5, which is likely involved in SA export from plastids (Rekhter et al., 2019), and mitochondrial arginase 2 (ARGAH2), which is highly induced by methyl jasmonate (Brownfield et al., 2008). Furthermore, the plastid Sigma1 interacting factor (Sib1, At3g56710) (Morikawa et al., 2002) is also highly expressed in *fmtclpc1* compared to *clpc1* (6.3-fold) and 2-3 fold compared to WT and *fmt*. Sib1 has been shown to be involved in JA and SA responses either within the chloroplast or in the nucleus (Xie et al., 2010; Zhang et al., 2022). Overexpression of Sib1 causes hyperactivation of defense-related genes following SA and JA treatments (Xie *et al*., 2010). These results suggest that JA and SA play an important role in the suppression of *clpc1* by *fmt*.

Some 30 transcription factors (TFs) have been highlighted as important during chloroplast and/or mitochondrial dysfunction, stress, retrograde signaling, or plastid biogenesis and proteostasis (Bittner *et al*., 2022; He et al., 2023; Hernandez-Verdeja, 2025; Wang et al., 2018; Wang *et al*., 2020) (Supplemental Table 3). Fifteen of these TFs showed at least one pairwise differential expression (adj. P<0.01) (Supplemental Data Set 1B; Fig. 4B). Four of these—HSFA2, GLK1, BBX15, and WRKY40 (At2g26150, At2g20570, At1g25440 and At1g80840)—showed significant differential expression patterns consistent with contributing to *clpc1* suppression (Fig. 4E). HSFA2 (cluster II-4) was strongly upregulated in *fmt* compared to WT but reduced in *clpc1,* with partial restoration in *fmtclpc1.* Notably, HSFA2 was also elevated in the *FMT/clpc1* background despite a more chlorotic and smaller phenotype, indicating that HSFA2 induction does not correlate with phenotypic recovery and is unlikely to be a primary driver of *clpc1* suppression. In contrast, GLK1 (cluster II-6) was unaffected in *fmt* but strongly suppressed in *clpc1* and *FMT/clpc1,* with partial restoration in *fmtclpc1.* The zinc-finger TF BBX15 (cluster II-4) (Ohmiya et al., 2019) was upregulated in *fmt* and downregulated in *clpc1* and *FMT/clpc1,* but restored to WT levels in *fmtclpc1*. WRKY40 (cluster III-9), which is involved in attenuation of basal pathogen defense and is upregulated by GLK1/2 (Lee et al., 2023), was elevated in *fmtclpc1,* likely reflecting a genetic interaction between *fmt* and *clpc1*. We note that the RNA levels of the well-studied TFs HY5 and GLK2 (both positive regulators of genes encoding chloroplast proteins involved in photosynthesis and chlorophyll synthesis), as well as the direct GLK1 target BBX14 (a homolog of BBX15) implicated in chloroplast development (Atanasov et al., 2024), were not significantly different (adj. P<0.01) across these five genotypes.

### Comparative proteomics reveals coordinated remodeling of chloroplast protein investment during *clpc1* suppression

To understand the suppression effect of *fmt* on the *clpc1* leaf proteome phenotype, total leaf proteomes of *clpc1-1* and *fmt-3clpc1-1* were compared by quantitative proteomics. A total of 2,975 proteins and protein groups comprising close homologs were identified, most of which (95%) were detected in both genotypes (Supplemental Data Set 2). Correlation analysis and principal component analysis (PCA) showed clear separation between genotypes and low variation among biological replicates (Supplemental Figure 5A,B). Of the identified proteins, 1,266 (43%) were localized to chloroplasts and/or mitochondria, representing ∼65% of all matched spectra. The five thylakoid complexes involved in the light reactions (PSII, PSI, cytochrome b6f, NDH and ATP synthase) all increased between 32% and 49%, whereas the Calvin-Benson cycle enzymes collectively increased by 25% in *fmtclpc1* compared with *clpc1*, consistent with suppression of the virescent phenotype (Supplemental Figure 5C). Consequently, relative investment in most other functional categories decreased, except for gluconeogenesis/glyoxylate cycle (bin 6), nitrogen metabolism (bin 12), and the TCA cycle/organic acid metabolism (bin 8), which increased by 9%, 4% and 3%, respectively. The relative abundance of the chloroplast ClpPRT complex was slightly lower (18%), whereas ClpC2 was 36% higher in *fmtclpc1* compared with *clpc1-1* (ClpC1 was not detected) (Fig. 5A), consistent with increased *CLPC2* RNA levels (Fig. 3D).

**Figure 5.**
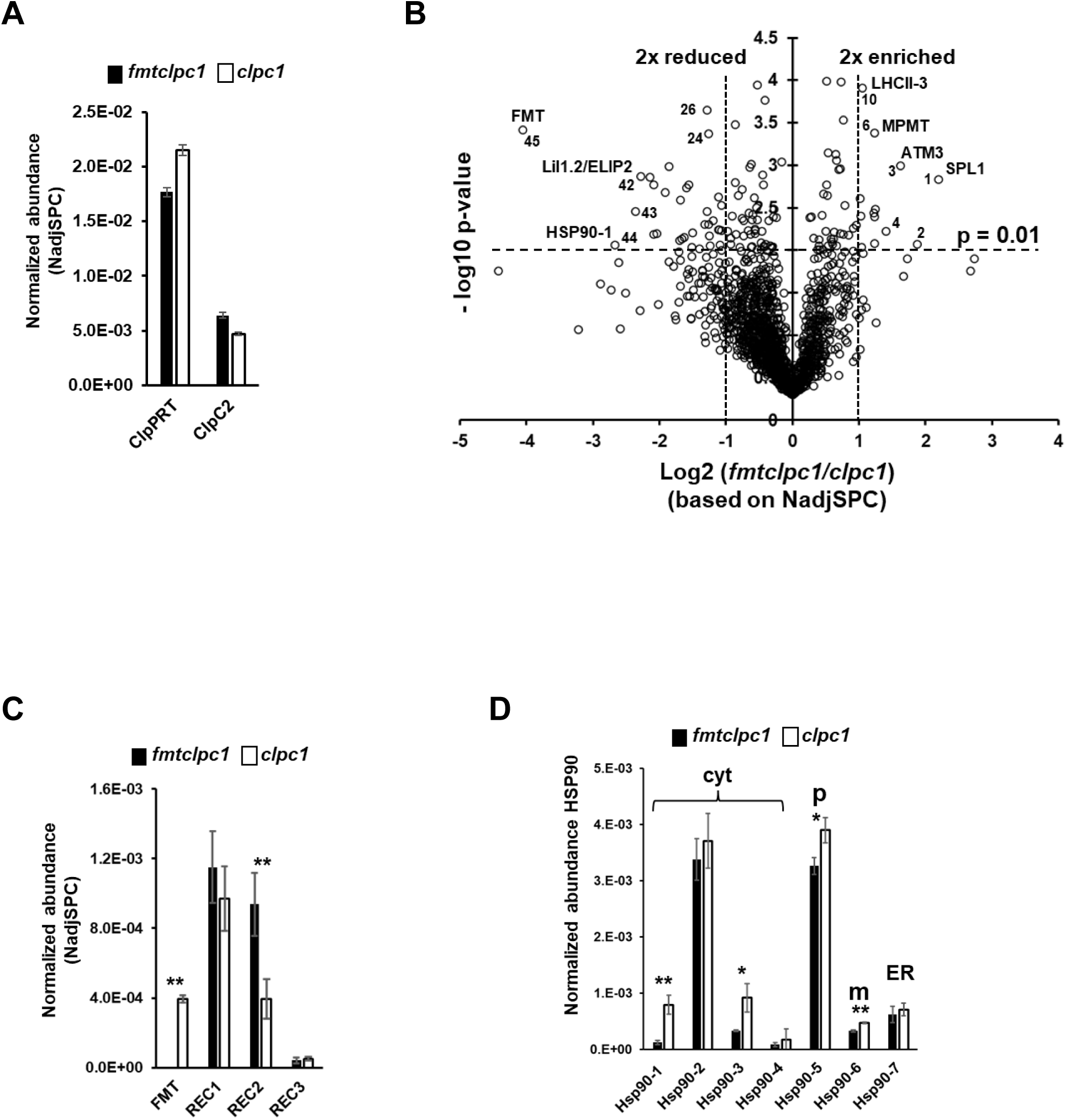
Comparative analysis of total rosette proteomes of *clpc1* and *fmtclpc1* soil- grown seedlings in growth stage 1.07 based on NadjSPC. **(A)** Bar diagram displaying the relative abundance of the chloroplast ClpPRT complex and ClpC2 in *fmtclpc1* and *clpc1*. **(B)** Volcano plot of the *fmtclpc1/clpc1* relative protein abundance ratio and significance of differential accumulation. Horizontal dashed lines indicate the -log10 p-value at 0.01 and the vertical dashed lines indicate 2-fold protein abundance difference between *fmtclpc1* and *clpc1*. **(C)** Bar diagram displaying the relative abundance of FMT and the related REC1,2,3 proteins in *fmtclpc1* and *clpc1*. *P<0.05; **P<0.01. **(D)** Bar diagram displaying the relative abundance of each of the seven HSP90 proteins in *fmtclpc1* and *clpc1*. *P<0.05; **P<0.01. HSP90-1,2,3,4 are localized to the cytosol. HSP90-5,6 and 7 are respectively localized to the chloroplast (p), mitochondria (m) and endoplasmic reticulum (ER).

Figure 5B shows the volcano plot of individual proteins comparing the two genotypes. At P<0.01 and a >2-fold difference between genotypes, 13 proteins increased and 32 decreased in *fmtclpc1* compared with *clpc1* (Fig. 5B, Table 1). All 13 increased proteins were plastid (8) or mitochondrial (4), except for REDUCED CHLOROPLAST COVERAGE 2 (REC2), which is localized to the cytosol. REC2 was enriched 2.3-fold in the double mutant. REC2 has two homologs (REC1 and REC3), and the REC family is phylogenetically related to FMT. REC1 and REC3 were detected in both genotypes, whereas FMT was detected only in *clpc1-1* and not in *fmtclpc1-1*, consistent with *fmt* being a null line; REC1 and REC3 did not change significantly (Fig. 5C).

**Table 1.**
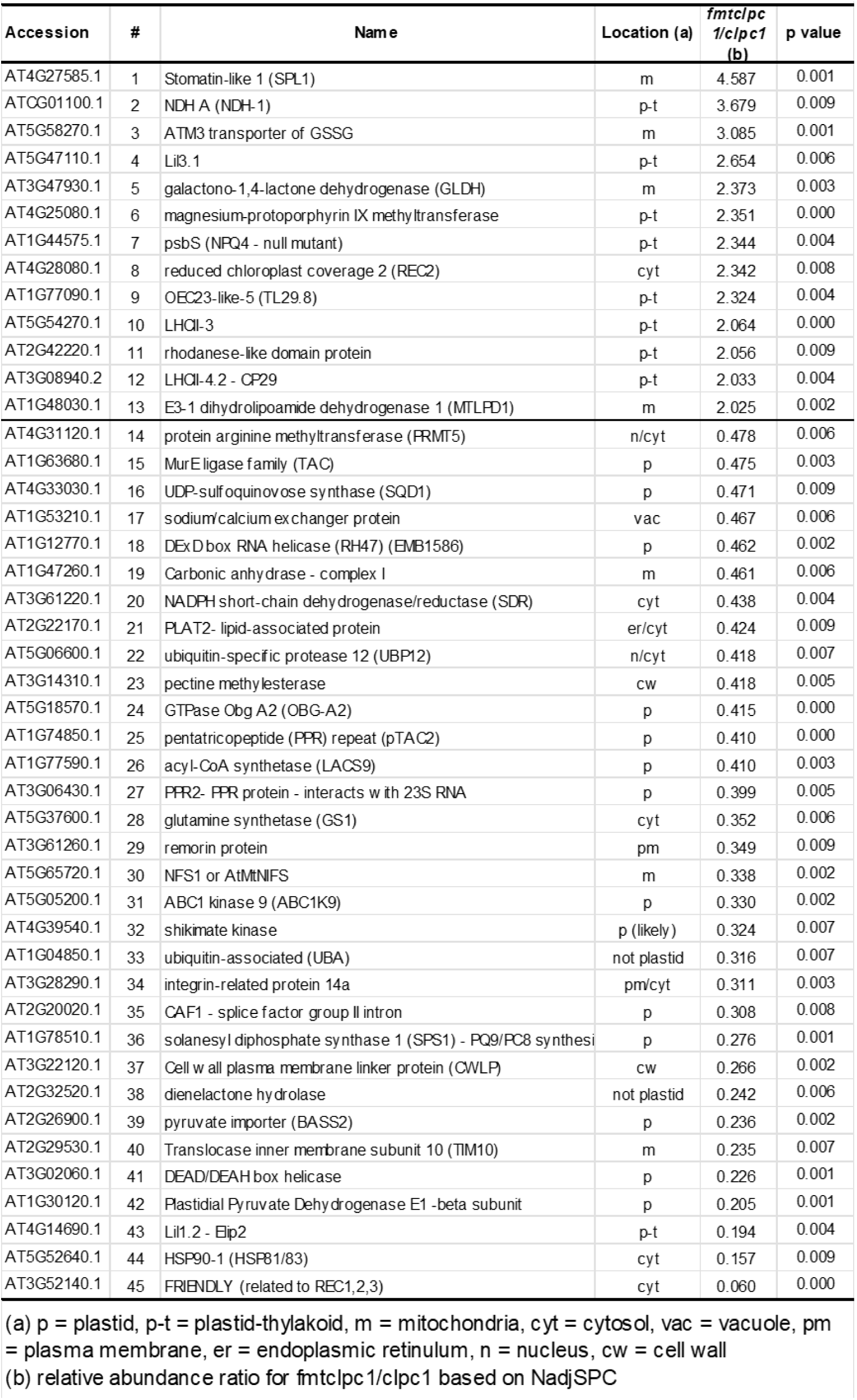
Differentially accumulating proteins (p<0.01; x<0.5 or x>2) based on NadjSPC in *clpc1* vs *fmtclpc1*.

Six of the increased chloroplast proteins are directly involved in the light reactions of photosynthesis, consistent with suppression of the virescent *clpc1* leaf phenotype. One additional chloroplast protein is involved in chlorophyll synthesis, whereas the thylakoid rhodanese domain protein has no known function. The mitochondrial protein showing the largest increase (4.6-fold) was stomatin-like protein 1 (SLP1), localized to the inner mitochondrial membrane (Fig. 5B, Table 1). SLP1, likely together with its homolog SLP2, forms a ∼3-MDa large complex in Arabidopsis mitochondria (Gehl *et al*., 2014). SLP2 was also strongly increased (6.7-fold) in *fmtclpc1* relative to *clpc1* (p-value 0.013). SLP1 and SLP2 are part of the band-7/SPFH superfamily of membrane scaffold proteins (Gehl and Sweetlove, 2014). Loss of SLP1 reduces protein abundance and activity of respiratory complex I, indicating a role in its assembly and/or stability (Gehl *et al*., 2014).

We also identified four mitochondrial prohibitins (PHB1, PHB2, PHB3, and PHB6), which are members of the same superfamily (Gehl and Sweetlove, 2014); however, their abundance was not increased in *fmtclpc1* compared with *clpc1*. Three additional mitochondrial proteins were increased: the membrane transporter ATM3, which exports oxidized glutathione; galactonolactone dehydrogenase (GLDH), which catalyzes the final step in ascorbate synthesis; and MTLPD1, a redox-regulated enzyme involved in photorespiration and the TCA cycle. These proteins are likely upregulated in response to mitochondrial dysfunction and stress resulting from loss of FMT. The ability of ATM3 to export GSSG has been implicated in helping to stabilize against excessive glutathione oxidation in mitochondria and thereby maintaining a suitable redox potential (Marty et al., 2019).

The 32 reduced proteins (*fmtclpc1* compared with *clpc1*) were distributed across chloroplasts (14), mitochondria (3), or other cellular compartments, encompassing diverse functions, particularly plastid fatty acid and lipid metabolism and plastid RNA-related processes (Table 1). The most strongly decreased proteins included FMT (as expected), cytosolic HSP90-1 involved in protein folding, the chloroplast thylakoid protein LIL1.2/ELIP2 (Tzvetkova-Chevolleau et al., 2007), the chloroplast pyruvate dehydrogenase E1-β subunit (PDH-Eβ) involved in acetyl-CoA synthesis for fatty acid biosynthesis, a chloroplast DEAD/DEAH-box helicase of unknown function, the essential mitochondrial intermembrane space protein TIM10 involved in protein sorting, and the chloroplast inner envelope pyruvate importer BASS2 (Furumoto et al., 2011). LIL1.2/ELIP2 reduces chlorophyll levels by inhibiting chlorophyll synthesis rather than increasing chlorophyll degradation, and LIL1.2/ELIP2 does not directly affect photosystem function or the electron transport chain (Tzvetkova-Chevolleau *et al*., 2007). Its low ratio (0.19) in *fmtclpc1/clpc1* is consistent with the low chlorophyll levels observed in *clpc1*. HSP90-1 is a cytosolic member of the large HSP90 family (7 members) of protein chaperones involved in protein folding, maturation, and degradation. HSP90-1 was reduced six-fold in *fmtclpc1* compared with *clpc1* (Fig. 5B, Table 1), indicating that cytosolic folding demand is higher in *clpc1* than in the double mutant. We identified all seven HSP90 members (HSP90-1 to -7), including chloroplast cpHSP90-5, mitochondrial HSP90-6 and ER-localized HSP90-7, as well as the four cytosolic members (HSP90-1 to -4) (Fig. 5D). All HSP90 members were lower in *fmtclpc1* than *clpc1,* indicating that suppression of *clpc1* resulted in reduced folding stress (or demands) across subcellular compartments (Wu et al., 2019).

### REC1 and REC2 are required for full suppression of the *clpc1-1* phenotype by *fmt*

In both our RNA-seq-based transcriptomic and MS/MS-based proteomic analyses, REC2 mRNA and protein were induced during *fmt*-mediated suppression of *clpc1-1*. Given its close phylogenetic relationship to *FMT*, we prioritized *REC2* together with its paralogs *REC1* and *REC3* for follow-up functional analysis. RT–qPCR (Fig. 6A) confirmed the RNA-seq results and was consistent with the proteomic analysis (Fig. 5C). *REC2* was increased 2.5-fold in *fmt-3clpc1-1*, and decreased to 0.6-fold in *FMT/clpc1-1* (OE#1), both relative to *clpc1-1* (Fig. 6A). In contrast, *REC1* and *REC3* transcript levels were largely unchanged, indicating that *REC2* is the only REC family member responsive to loss of *FMT* function.

**Figure 6.**
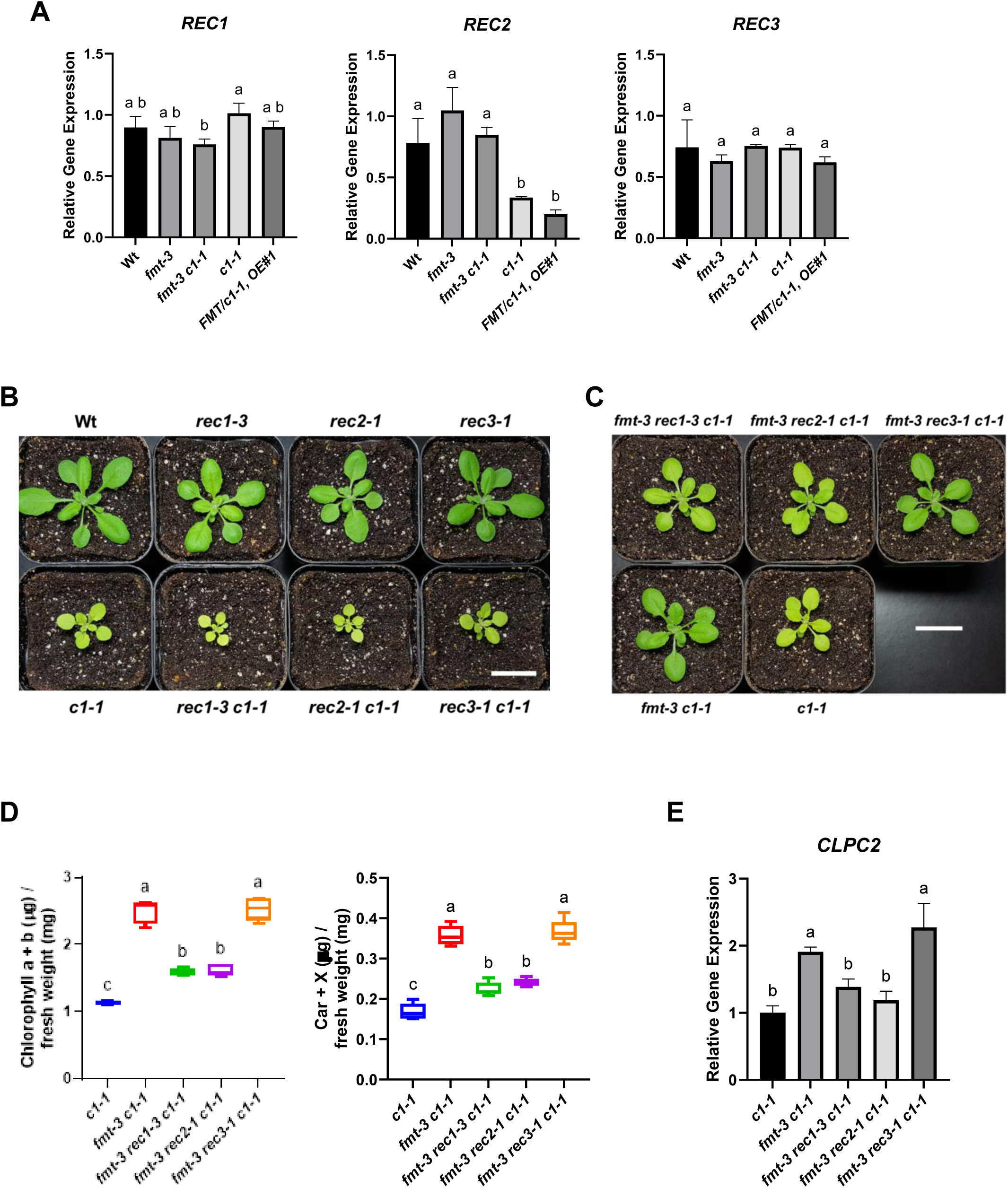
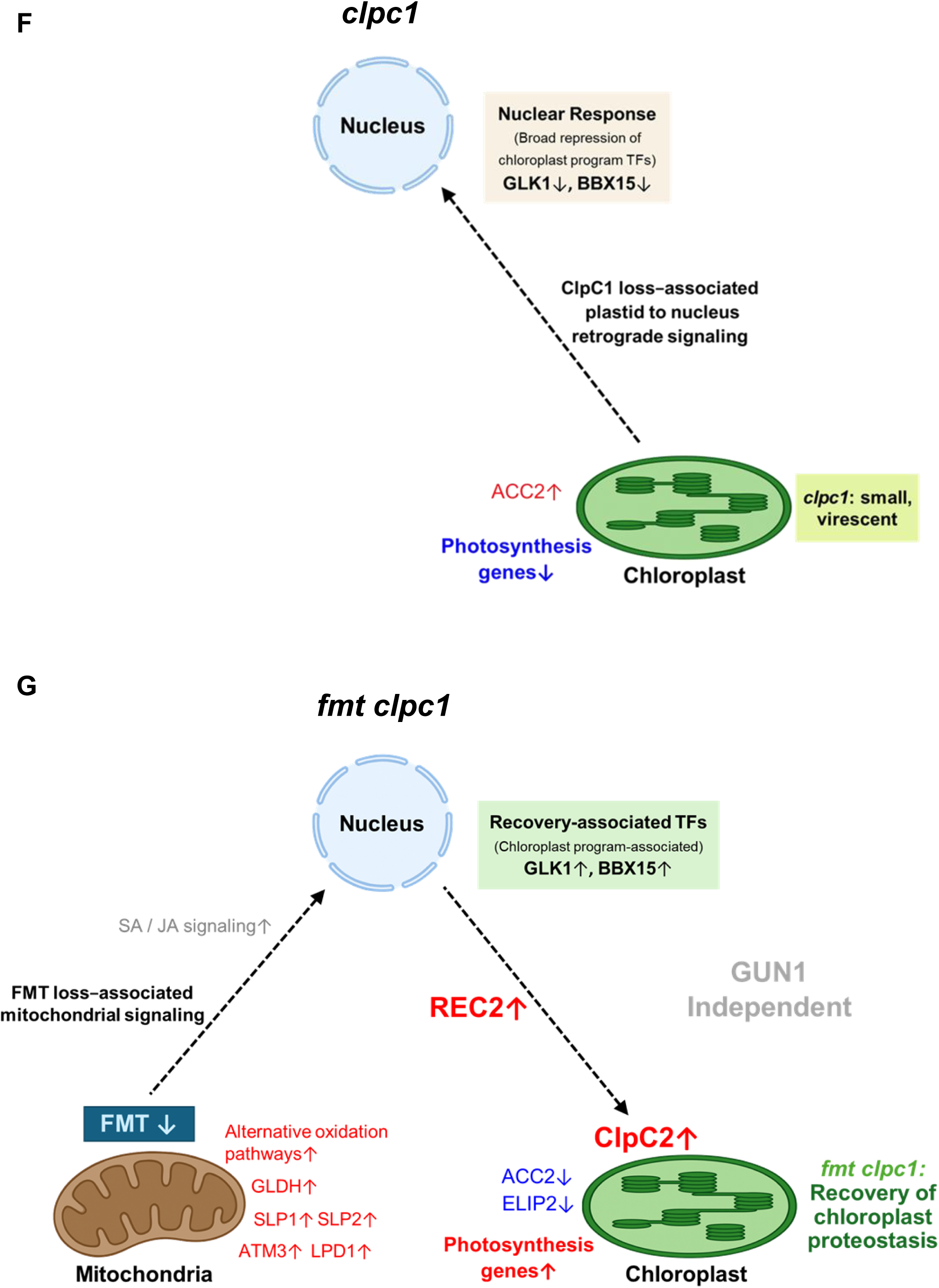
*REC1* and *REC2* are required for full suppression of *clpc1-1* by *fmt*. **(A)** Quantitative RT-PCR analysis of *REC1*, *REC2*, and *REC3* transcript levels in wild type (Col-0), *fmt-3*, *fmt-3 clpc1-1*, *clpc1-1*, and *FMT/clpc1-1* OE#1. Data represent mean ± SD from three biological replicates. **(B)** Growth and development of wild type, *clpc1-1*, *rec1-3*, *rec2-1*, *rec3-1*, *rec1-3 clpc1-1*, *rec2-1 clpc1-1*, and *rec3-1 clpc1-1* mutants. Plants were grown on soil for 17 days under a 16/8 hour light/dark cycle at 100 μmol photons m⁻² s⁻¹. Scale bar = 2 cm. **(C)** Growth and development of *fmt-3 rec1-3 clpc1-1*, *fmt-3 rec2-1 clpc1-1*, *fmt-3 rec3-1 clpc1-1*, *fmt-3 clpc1-1*, and *clpc1-1* mutants. Plants were grown on soil for 18 days under a 16/8 hour light/dark cycle at 100 μmol photons m⁻² s⁻¹. Scale bar = 2 cm. **(D)** Pigment accumulation in the same lines shown in (C). Total chlorophyll and carotenoid contents were measured in 21-day-old plants grown under a 16/8 hour light/dark cycle at 100 μmol photons m⁻² s⁻¹. Total chlorophyll (left) and total carotenoids (right) are shown. Pigment contents are expressed as micrograms per milligram fresh weight (μg/mg FW). Values represent mean ± SD (n = 5). **(E)** Quantitative RT-PCR analysis of *CLPC2* transcript levels were measured in *clpc1-1*, *fmt-3 clpc1-1*, *fmt-3 rec1-3 clpc1-1*, *fmt-3 rec2-1 clpc1-1*, and *fmt-3 rec3-1 clpc1-1*. Data represent mean ± SD from three biological replicates. (A), (D), and (E) Different lowercase letters indicate groups that differ significantly, as determined by one-way ANOVA followed by Tukey’s range test (P < 0.05). **(F) *clpc1*-associated plastid stress state.** Working model illustrating the transcriptional and physiological state of *clpc1* relative to wild type. Loss of ClpC1 causes chloroplast dysfunction, triggering plastid-to-nucleus retrograde signaling associated with a broadly repressive nuclear response. This state correlates with reduced expression of multiple transcription factors, including GLK1 and BBX15, as well as decreased expression of photosynthesis-related genes. In parallel, plastid-localized metabolic response genes such as *ACC2* are upregulated, reflecting altered plastid metabolic states associated with impaired chloroplast proteostasis and the small, virescent *clpc1* phenotype. Dashed arrows denote inferred signaling relationships based on transcriptomic and phenotypic analyses. **(G) FMT-mediated recovery state in *fmtclpc1*.** Working Model illustrating the transcriptional and physiological state associated with FMT-dependent suppression of *clpc1*. Loss of *FMT*, a mitochondria-associated member of the FMT–REC gene family, is associated with mitochondrial perturbation that engages a mitochondria-to-nucleus communication route. This nuclear response correlates with increased expression of *REC2* and *CLPC2*, the latter encoding a plastid Clp chaperone that compensates for loss of ClpC1 and promotes restoration of chloroplast proteostasis in *fmtclpc1*. In this recovery state, transcription factors (TFs) associated with chloroplast gene programs (GLK1, BBX15) are upregulated, photosynthesis-related genes are reactivated, and plastid stress markers such as *ACC2* and *ELIP2* are reduced. Upregulation of mitochondrial redox- and metabolism-associated factors (e.g., SLP1, SLP2, ATM3, GLDH, LPD1) is consistent with mitochondria-derived signaling. Increased expression of salicylic acid (SA)- and jasmonic acid (JA)-associated signaling components further reflects a remodeled nuclear transcriptional state. The *fmt–clpc1* interaction occurs independently of GUN1, which is depicted separately. Genes shown in red are upregulated and those in blue are downregulated based on integrated transcriptomic and proteomic datasets; dashed arrows denote inferred signaling or transcriptional relationships supported by genetic, transcriptomic, and proteomic evidence.

We next tested whether *REC1*, *REC2*, or *REC3* might share *FMT*’s capacity to suppress the *clpc1-1* phenotype. To address this, we generated double mutants between *clpc1-1* and *rec1-3*, *rec2-1*, or *rec3-1*. As shown in Figure 6B, none of these double mutants exhibited suppression of the *clpc1-1* phenotype. The *rec3-1clpc1-1* mutant closely resembled *clpc1-1* alone, whereas both *rec1-3clpc1-1* and *rec2-1clpc1-1* displayed smaller rosettes and more pronounced chlorosis than *clpc1-1*. These results indicate that, although evolutionarily related to *FMT*, the *REC* genes cannot substitute for *FMT* function in mitigating *CLPC1* deficiency.

We next tested whether *REC* genes are required for *fmt*-mediated rescue. Both *fmt-3rec1-3clpc1-1* and *fmt-3rec2-1clpc1-1* exhibited intermediate phenotypes between *fmt-3clpc1-1* and *clpc1-1*, whereas *fmt-3rec3-1clpc1-1* resembled *fmt-3clpc1-1* (Fig. 6C). Pigment analyses mirrored these patterns: total chlorophyll and carotenoids were reduced in *fmt-3rec1-3clpc1-1* and *fmt-3rec2-1clpc1-1* relative to *fmt-3clpc1-1*, while *fmt-3rec3-1clpc1-1* remained unchanged (Fig. 6D). *CLPC2* transcript abundance followed the same trend—reduced in *fmt-3rec1-3clpc1-1* and *fmt-3rec2-1clpc1-1*, unchanged in *fmt-3rec3-1clpc1-1* (Fig. 6E). Together, these results show that *REC1* and *REC2* are required for full *fmt*-mediated suppression of *clpc1-1*, whereas *REC3* is dispensable. *REC1* appears to be constitutively expressed, while *REC2* is upregulated in response to the loss of *FMT*.

## DISCUSSION

Our study reveals an unexpected connection between mitochondrial dysfunction and chloroplast proteostasis, showing that loss of *FMT–*a regulator of mitochondrial morphology (El Zawily *et al*., 2014; Logan *et al*., 2003)—enables a compensatory response that restores chloroplast function in the absence of ClpC1. Although *fmt* mutants exhibit clustered mitochondria and impaired mitochondrial dynamics, they paradoxically display improved chloroplast performance in the *clpc1* background. These findings indicate that mitochondrial perturbation in *fmt* does not simply exacerbate cellular stress but is associated with selective transcriptional and proteomic changes—most notably induction of *CLPC2*—that enable functional compensation for loss of ClpC1. A working model summarizing these contrasting cellular states is presented in Fig. 6F,G.

In the *clpc1* mutant (Fig. 6F), loss of ClpC1 establishes a plastid stress state associated with plastid-to-nucleus retrograde signaling and a broadly repressive nuclear response. This state correlates with reduced expression of transcription factors such as GLK1 and BBX15 and decreased expression of photosynthesis-related genes. In parallel, plastid-localized metabolic response genes including *ACC2* are upregulated, reflecting altered plastid metabolic status linked to impaired chloroplast proteostasis and the small, virescent phenotype. Together, these features define a stress-associated transcriptional configuration that is insufficient to restore plastid function.

By contrast, loss of FMT shifts the system into a recovery state (Fig. 6G). Despite persistent mitochondrial clustering, *fmt*-mediated suppression restores chloroplast ultrastructure, photosynthetic protein accumulation, and Clp-dependent proteolysis. This transition correlates with induction of *REC2* and derepression of *CLPC2*, enabling ClpC2 to functionally substitute for ClpC1. In this recovery state, transcription factors associated with chloroplast gene programs, including GLK1 and BBX15, are restored, photosynthesis-related genes are reactivated, and plastid stress markers such as *ACC2* and *ELIP2* decline. These changes occur independently of GUN1-dependent retrograde signaling, indicating engagement of a distinct mitochondria-to-nucleus communication route rather than canonical plastid stress pathways. Although GUN1 is known to buffer import-associated defects (Wu *et al*., 2019) and to stabilize specific plastid RNAs to maintain organellar gene expression (Tang *et al*., 2024), these responses alone are insufficient to restore chloroplast function in *clpc1*. The ability of *fmt* to suppress *clpc1* in the absence of GUN1, together with the persistence of mitochondrial abnormalities in suppressed lines, indicates that mitochondrial perturbation promotes plastid recovery through a mechanism distinct from established plastid-to-nucleus signaling pathways.

The recurrence and strength of *fmt* alleles among independent suppressors highlight a specific role for FMT in enabling compensation for ClpC1 loss. Although additional suppressor lines were recovered in the screen, none displayed the near-complete phenotypic restoration observed in *fmt* mutants, underscoring the distinctive contribution of *FMT*. Loss of *FMT* does not mitigate defects in other components of the Clp protease system (Kim *et al*., 2015; Rudella *et al*., 2006) or chloroplast protein import (Bedard *et al*., 2017; Kubis *et al*., 2003), further emphasizing that the compensatory response is not a general bypass of plastid dysfunction.

A key downstream effector of this compensatory transition is *CLPC2*, a ClpC1 paralog. While ClpC2 plays a minor role under normal conditions (Kovacheva *et al*., 2007; Nishimura *et al*., 2013) and has been implicated in responses to organelle perturbation (Lopez *et al*., 2024), our data extend this view by showing that mitochondrial perturbation is sufficient to drive *CLPC2* transcription to levels that restore Clp-dependent proteolysis and chloroplast biogenesis. The proportional increases in *CLPC2* expression observed in native-promoter overexpression lines indicate that the promoter is not saturated in *clpc1-1*, suggesting that *fmt* derepresses *CLPC2*. Conversely, increasing *FMT* abundance reduces *CLPC2* expression and produces phenotypes resembling the *clpc1-1clpc2-2* double mutant, consistent with an indirect, dose-dependent negative influence of *FMT* on *CLPC2* expression.

The integrated RNA-seq and comparative proteomics analyses provide critical context for this genetic compensation. Rather than revealing activation of a dominant stress or retrograde signaling pathway, these datasets show coordinated remodeling of nuclear gene expression and chloroplast protein investment in *fmtclpc1*. Transcriptome profiling indicates partial restoration of photosynthesis-associated gene expression and attenuation of plastid proteotoxic stress signatures, accompanied by selective remodeling of stress- and signaling-associated gene clusters. Notably, Cluster III highlights genes involved in hormonal (JA- and SA-related) and metabolic adjustment that are selectively enriched in *fmtclpc1*, supporting the idea that suppression involves entry into a distinct adaptive cellular state rather than engagement of a linear signaling cascade. Comparative proteomics reinforces this interpretation by demonstrating recovery of chloroplast protein accumulation and reduced cytosolic chaperone burden, consistent with improved plastid function rather than a generalized stress response.

Genetic and proteomic analyses further implicate REC proteins in linking mitochondrial perturbation to *CLPC2* induction. *REC2* is strongly enriched in *fmtclpc1*, and both *REC1* and *REC2* are required for full *fmt*-mediated suppression, whereas *REC3* appears dispensable.

Together with previous studies showing roles for REC proteins in plastid compartment size (Hu *et al*., 2024; Larkin *et al*., 2016) and plastid-related gene expression (Hu *et al*., 2024), our findings support a model in which *REC1* and *REC2* act genetically upstream of *CLPC2* induction and contribute to the broader remodeling of plastid proteostasis during the transition from the *clpc1* stress state to the *fmt*-mediated recovery state.

Previous work has identified GOLDEN2-LIKE and B-BOX transcription factors as components of transcriptional networks linking chloroplast functional status with nuclear gene regulation (Susila et al., 2023). In our study, *GLK1* and *BBX15* showed genotype-dependent expression changes across *clpc1*, *fmt*, and *fmtclpc1*, consistent with their placement within the two-state model outlined in Fig. 6F,G. By contrast, GLK2, BBX14, and BBX16 did not exhibit coordinated expression changes, suggesting context-dependent engagement of the GLK–BBX regulatory network.

Together, these findings support a model in which mitochondrial perturbation exposes a latent capacity for inter-organelle compensation, enabling selective transcriptional and proteomic remodeling that restores chloroplast proteostasis when ClpC1 function is compromised. Rather than simply alleviating stress, loss of FMT appears to shift the cellular signaling landscape from a repressive plastid stress state to a compensatory recovery state.

## MATERIALS AND METHODS

### EMS Mutagenesis and Suppressor Mapping by Whole-Genome Sequencing

Approximately 10,000 *Arabidopsis thaliana clpc1-1* seeds (Col-0 background) were subjected to ethyl methanesulfonate (EMS) mutagenesis following standard procedures (Kim et al., 2006). Treated seeds (M1 generation) were sown on soil and grown under long-day conditions (16/8 hour light/dark cycle at 100 μmol photons m⁻² s⁻¹, 22 °C). M2 seeds were collected in bulk from individual M1 plants and screened for suppressor phenotypes. The *clpc1-1* mutant exhibits a strong chlorotic and stunted phenotype, allowing visual identification of M2 seedlings with restored leaf greening and improved growth. Approximately 60 groups of M2 seedlings were examined, and ∼10 independent suppressor candidates showing partial or near-complete phenotypic recovery were identified. Four lines exhibiting the strongest recovery were selected for further analysis and designated *socc* (*<u>s</u>uppressor <u>o</u>f <u>c</u>lp<u>c</u>1*). Each *socc* line was backcrossed to the parental *clpc1-1* mutant to generate BC1F2 populations. Segregation of the suppressor phenotype among BC1F2 progeny followed an approximate 1:3 ratio (green:chlorotic), consistent with monogenic recessive inheritance.

For bulked segregant analysis, leaf tissue from at least 50 green and 50 chlorotic BC1F2 plants was harvested separately. Genomic DNA was extracted using the DNeasy Plant Mini Kit (Qiagen), and DNA from plants of the same phenotype was pooled to generate “green” and “chlorotic” bulks. Whole-genome sequencing (WGS) was performed by BGI Genomics (Hong Kong). Candidate causal variants were identified by comparing allele frequency differences between the green and chlorotic bulks using the SIMPLE pipeline (Wachsman *et al*., 2017). Putative causal mutations were further validated by genetic complementation.

### Plant Growth, T-DNA Mutant Isolation, and RNA Preparation and RT-PCR Analysis

Plant growth, genotyping, and RNA extraction were performed as described previously (Kim *et al*., 2009). Various growth conditions are detailed in the figure legends. All T-DNA insertion lines were in the Columbia background and are as follows: *FMT* (AT3G52140), SALK_046271 (*fmt-3*) and SALK_056717 (*fmt-4*); *CLPC1* (AT5G50920), SALK_014058 (*clpc1-1*); *CLPC2* (AT3G48870), SAIL_622_B05 (*clpc2-2*); *CLPR2* (AT1G12410), SALK_046378 (*clpr2-1*); *CLPT1* (AT4G25370), GK-285A05 (*clpt1-2*); *CLPT2* (AT4G12060), SAIL_340A10 (*clpt2-1*); *TOC33* (AT1G02280), N2107726 (*ppi1-1*); *TIC40* (AT5G16620), SAIL_92_C10 (*tic40-4*); *GUN1* (AT2G31400), SAIL_33_D01 (*gun1-101*); *REC1* (AT1G01320), SALK_049453C (*rec1-3*); *REC2* (AT4G28080), SALK_020337C (*rec2-1*); and *REC3* (AT1G15290), SAIL_1164_H02 (*rec3-1*).

For RNA-seq analysis, total RNA was isolated using the RNeasy Plant Mini Kit (Qiagen). For RT-PCR, total RNA was isolated using either the RNeasy Plant Mini Kit (Qiagen) or the XENOPURE Total RNA Purification Kit (CelltoBio). First-strand cDNA was synthesized from equal amounts of total RNA using SuperScript III Reverse Transcriptase (Invitrogen) or TOPscript Reverse Transcriptase (Enzynomics). Semi-quantitative RT-PCR and RT-qPCR were performed as described previously (Bae et al., 2025; Kim *et al*., 2015). Graphs were generated using GraphPad Prism 10. Primers used for genomic PCR, RT-PCR analysis, and various complementation assays are listed in table S2.

### Cloning and Complementation

For *FMT*, the full-length native promoter and 5′UTR of *FMT* (1.0 kb), together with the full-length *FMT* genomic DNA (8.0 kb), totaling 9.0 kb, were PCR-amplified using KOD polymerase (TOYOBO). The PCR products were cloned into a modified pCR8/GW/TOPO vector (Invitrogen), into which BamHI and NotI restriction sites were introduced. Using LR Clonase (Invitrogen), the DNA fragment was transferred into the pGWB4 Gateway destination plant binary vector. For *CLPC2*, the full-length native promoter and 5′UTR of *CLPC2* (1.9 kb), together with the full-length *CLPC2* genomic DNA (5.6 kb), totaling 7.5 kb, were PCR-amplified using KOD polymerase (TOYOBO). The PCR products were cloned into a modified pCR8/GW/TOPO vector (Invitrogen), into which PstI and NotI restriction sites, as well as a StrepII tag and a *nosT* sequence (0.5 kb), were introduced. Using LR Clonase (Invitrogen), the DNA fragment was transferred into the pMDC99 Gateway destination plant binary vector. The primers used are listed in table S2. Competent cells of *Agrobacterium tumefaciens* strain GV3101 were transformed with the binary vectors. Plant transformation and selection were performed as described previously (Kim *et al*., 2009), using the *Agrobacterium*-mediated floral dip method with selection on ½ MS agar plates containing 25 μg/mL hygromycin (Duchefa Biochemie). Complemented lines were identified and verified by PCR genotyping.

### Generation of Mutant Lines Using CRISPR/Cas9

Mutations in *CLPC1* were generated using the CRISPR/Cas9 system. sgRNAs were designed with CRISPR-P v2.0 and cloned into the pBR-Cas9 vector (provided by Dr. Jae-Young Yun, SeoulTech, Seoul, Korea). *Arabidopsis thaliana* Col-0 plants were transformed by the Agrobacterium-mediated floral dip method (*A. tumefaciens* GV3101). Transgenic plants were screened by PCR amplification of the target region followed by Sanger sequencing of CTAB-extracted DNA. Two independent knockout alleles were recovered: *clpc1-4*, carrying a 1-bp insertion in exon 4, and *clpc1-5*, containing a 32-bp deletion plus a single-nucleotide substitution in the same exon. Both mutations introduce frameshifts predicted to abolish ClpC1 protein function. Cas9-free homozygous segregants were identified by PCR-based genotyping (primer sequences in table S2) and used for all subsequent analyses.

### Pigment Analysis

Chlorophyll and carotenoid contents on a fresh-weight basis were measured in 80% acetone as described previously (Kim *et al*., 2013).

### Photosynthetic Efficiency Analysis

Fv/Fm measurements were performed as described previously (Bae *et al*., 2025). Briefly, plants were dark-adapted for 30 min at room temperature, and chlorophyll fluorescence parameters (Fo and Fm) were recorded using a FluorCam 800MF (Photon Systems Instruments, Czech Republic). Fv/Fm values were calculated as (Fm - Fo)/Fm to evaluate photosystem II photochemical efficiency under normal growth conditions.

### Transmission Electron Microscopy (TEM)

Leaves from 23-day-old plants were fixed with a mixture of 2% (v/v) glutaraldehyde and 2% (v/v) paraformaldehyde in 0.05 M cacodylate buffer (pH 7.2) at room temperature for 4 h. The samples were washed with the same buffer and post-fixed with 1% (w/v) OsO₄ in 0.05 M cacodylate buffer at room temperature for 1 h. Fixed samples were washed again with the buffer and dehydrated through a graded ethanol series (30–100%). The samples were then embedded in LR White resin at 50 °C for 24 h. Ultrathin sections (80–100 nm thick) were prepared using an ultramicrotome equipped with a diamond knife. Sections were stained with uranyl acetate and lead citrate and examined using a transmission electron microscope (JEM-2100F, JEOL Ltd., Japan).

### Chloroplast Isolation and Total Leaf Protein Extraction

For chloroplast isolation, leaves from wild-type and various mutant alleles were briefly homogenized in grinding medium (50 mM HEPES-KOH, pH 8.0, 330 mM sorbitol, 2 mM EDTA, 5 mM ascorbic acid, 5 mM cysteine, and 0.03% BSA) and filtered through a nylon mesh. The crude plastids were collected by centrifugation at 1,100 g for 2 min and further purified on 35–85% Percoll cushions (Percoll in 0.6% Ficoll and 1.8% polyethylene glycol) by centrifugation at 3,750 g for 10 min, followed by one additional wash in grinding medium lacking ascorbic acid, cysteine, and BSA. Chloroplasts were subsequently lysed in 10 mM HEPES-KOH, pH 8.0, 5 mM MgCl₂, and 15% glycerol, containing a mixture of protease inhibitors, under mild mechanical disruption. Protein concentrations were determined using the BCA Protein Assay Kit (Thermo Scientific). Total leaf proteins were extracted as described previously (Kim *et al*., 2015).

### Immunoblot Analysis

For immunoblotting, proteins were transferred onto nitrocellulose or polyvinylidene fluoride (PVDF) membranes and probed with specific antibodies using chemiluminescence detection, following standard procedures. The antisera against *Arabidopsis* ClpF and ClpS1 were described previously (Nishimura *et al*., 2015). The anti-PAA2 antibody was kindly provided by Dr. Marinus Pilon (Colorado State Univ., Fort Collins, CO, USA).

### PAA2 Degradation Assay

The PAA2 degradation assay was performed as described previously (Tapken *et al*., 2015) with minor modifications. Briefly, *Arabidopsis thaliana* seedlings were grown on half-strength MS agar plates (pH 5.8) containing 0.6% (w/v) agar and 1% (w/v) sucrose for 21 days under short-day conditions (10/14 hour light/dark cycle at 100 μmol photons m⁻² s⁻¹). For copper treatments, 5 μM CuSO₄ was added to the growth medium, and for control conditions, 0.05 μM CuSO₄ was used. Total leaf protein extraction and immunoblot analysis were performed as described previously (Kim *et al*., 2015). PAA2 abundance was detected using affinity-purified anti-PAA2 antibodies (1:1000 dilution).

### RNAseq analysis

RNA analysis was carried out for five genotypes of Arabidopsis seedlings in stage 1.07, with 3 biological replicates. The genotypes are: WT, *clpc1-1*, *fmt-3*, *fmt-3 clpc1-1*, FMT overexpressed line OE#1 in *clpc1-1* (*FMT/clpc1*). Total RNA was isolated with an RNeasy plant mini kit (Qiagen). Total RNA samples were processed for library preparation and sequencing following the manufacturer’s protocol. Briefly, mRNA was enriched from total RNA using oligo(dT) beads targeting the poly(A) tails. The isolated mRNA was then fragmented into smaller pieces, which served as templates for first-strand cDNA synthesis using random primers. Subsequently, second-strand cDNA synthesis was performed, incorporating dUTP in place of dTTP to enable strand specificity. The resulting double-stranded cDNA underwent end-repair and 3′-adenylation, after which sequencing adapters were ligated. The adapter-ligated fragments were then amplified via PCR, and the amplified libraries were subjected to quality control assessment. Following quality control, the libraries were circularized and further amplified to form DNA nanoballs (DNBs). Finally, sequencing was performed using the DNBSEQ platform (DNBSEQ Technology). Raw data with adapter sequences or low-quality sequences was filtered using SOAPnuke (Chen et al., 2018). For individual replicate-level expression analyses, the HISAT2 aligned bam files were used for reads counting using the featureCounts program (Liao et al., 2014)

Data normalization (Supplemental Data Set 1A), downstream analysis of differentially expressed genes, and adjusted p-value calculation were performed using the DESeq2 package v.1.44.0 (Love et al., 2014) in R v4.4.3. . Specifically, a DESeqDataSet object was generated using the raw counts, with design = ∼genotype. Pairwise results for each genotypic comparison of interest were then generated using the ‘results()’ function of DESeq2, with the function’s contrast parameter being used to specify the relevant genotypes. The p-values were determined by the default Wald test, and these p-values were adjusted using the Benjamini-Hochberg method. Genes with a differential expression of at least two-fold (x<0.5 or x>2) and adjusted p value <0.01 in one of the 10 possible comparisons across genotypes were considered for further investigation. This resulted in a set of 4745 nuclear protein-coding genes that we annotated for subcellular protein location (in particular chloroplast and mitochondria) and function (using the MapMan bin system) from public resources (TAIR, PPDB and publications) (Supplemental Data Set 1B). The gene clustering was generated using the Ward D2 method with Euclidean distance in the pheatmap v1.0.13 R package (Supplemental Data Set 1B). PCA analysis was performed using the GLEE software (Kim *et al*., 2013). The graphs are generated either using Microsoft Excel or GraphPad Prism 10.

### Comparative Proteome Analysis

Total proteome extracts of seedlings of *clpc1-1* and *fmt-3 clpc1-1* (each with three biological replicates) were separated by Tris-Glycine SDS-PAGE using 8-16% precast gels (Cat. No. KG7550-BC; KOMA Biotech, Korea). Gels were stained with Coomassie Brilliant Blue. Each of the six SDS-PAGE gel lanes was completely cut into consecutive gel slices (10 slices per lane), followed by reduction, alkylation, and in-gel digestion with trypsin as described in (Friso et al., 2011). The extracted peptides were resuspended in 2% formic acid and analyzed using a QExactive mass spectrometer equipped with a nanospray flex ion source and interfaced with a nanoLC system and autosampler (Dionex Ultimate 3000 Binary RSLCnano system) as described in (Rei Liao *et al*., 2022) with the exception that AGC target values were set at 1 × 10^6^ for the MS survey scans and maximum scan time 30 ms and 5 × 10^5^ for MSMS scans and maximum scan time 50 ms [Other parameters: MS: 70k resolution, scan range 400-2000 m/z; MSMS: 17.5k resolution, top10, loop count 10, isolation window 2.0 m/z, dynamic exclusion 15 sec].

Peak lists in MGF format were generated from RAW files using Distiller software (version 2.7.1.0) in default mode (Matrix Science). MGF files were searched with MASCOT v2.4.0 against TAIR10 including a set of typical contaminants and the decoy (71148 sequences; 29099536 residues). Two parallel searches (Mascot p-value <0.01 for individual ion scores; precursor ion window 700 to 3500 Da) were carried out: (i) Full tryptic (error tolerance 10 ppm for MS and 0.5 Da for MS/MS) and (ii) Semi-tryptic (error tolerance 6 ppm and 0.5 Da for MS/MS) in both cases with fixed Cys-carbamido-methylation and variable M-oxidation, N-terminal acetylation, N-terminal formylation and 2 missed cleavages (in Mascot PR or PK does not count as missed cleavage). In-house post-Mascot filtering removed redundancies between the full and semi-tryptic searches and removed spectral matches with ion scores below 33. Proteins could only be identified by the spectral counting method (SPC) with the full tryptic (6 ppm) search. The semi-tryptic search served to increase protein coverage and was combined with the full tryptic search results. Final false discovery rate for proteins identified with one spectral match was zero for those with two or more spectral matches. Proteins identified by MS/MS spectra that were all shared with other proteins identified by unique peptides were discarded.

Proteins were quantified by the spectral counting method (SPC) using full and semi-tryptic peptides search results. For quantification by spectral counting, each accession was scored for total spectral counts (SPC), unique SPC (uniquely matching to an accession) and adjusted SPC (Friso *et al*., 2011). The latter assigns shared peptides to accessions in proportion to their relative abundance using unique spectral counts for each accession as a basis. Proteins that shared more than 80% of their matched peptides with other proteins across the complete dataset were grouped into families quantified as groups with these homologs (Friso *et al*., 2011). After selecting only the best scoring isoform model per protein, proteins that shared a significant number of matched spectra were grouped, and proteins and groups with less than 3 adjSPC were discarded for further analysis; this resulted in 2976 proteins including 124 groups. For significance analysis and Volcano plots to compare the two genotypes, the minimal total adjSPC was raised to >12, resulting in 1529 proteins and groups; this ensured a more robust dataset and lower FDR. Within each sample, a value of 0.99 AdjSPC was imputed for all proteins with missing values. This was followed by normalization of the adjSPC to the total adjSPC of the samples resulting in NadjSPC for each protein. Average NadjSPC per genotype across the 3 replicates were calculated and *fmtclpc1/clpc1* NadjSPC ratios were determined for each protein. p-value and q-value (FDR) were then calculated to determine proteins that showed significant abundance differences between *clpc1* and *fmtclpc1*. We created Volcano plots and plotted total protein abundance against the protein abundance ratios.

## Supporting information

Supplemental Data Set 1

Supplemental Data Set 2

## ACKNOWLEDGMENTS

This research was supported by the following grants: National Research Foundation of Korea (2019R1I1A1A01064009 and RS-2023-00247582 to J.K., RS-2023-00208784 to D.W.L., RS-2024-00407469 and RS-2025-00564152 to B.h.L., and NRF-2022R1A3B1078180 to J.H.A.), and a US National Science Foundation grant MCB-2322813 to K.J.v.W.

## AUTHOR CONTRIBUTIONS

J.K. conceived the experiments, and K.J.v.W. conceived the multi-omics experiments. J.K. performed EMS screening, all genetic crosses, and data analysis. K.J.v.W. and P.R. performed quantitative comparative proteomics and RNA-seq analyses, while Z.N. conducted the initial RNA-seq analysis. N.S. and H.L. provided bioinformatics consultation and support. J.K., C.N., and D.B.K. performed biochemical experiments, including total and chloroplast protein extraction and immunoblotting. J.K., D.B.K., and J.-Y.K. performed cloning. C.N. carried out the PAA2 degradation assay, and J.-Y.K. generated CRISPR/Cas9 lines. J.K., N.B., and H.K. performed RT-PCR analyses. J.K. prepared samples for RNA-seq and proteomics. J.K., C.N., N.B., H.K., D.B.K., R.L., J.H.K., and G.C. performed plant transformation and/or other molecular genetic experiments. J.K., J.H.A., B.h.L., D.W.L., and K.J.v.W. supervised the project and analyzed and interpreted the data. J.K. and K.J.v.W. wrote the manuscript. D.W.L. and P.R. reviewed, edited, and provided substantive feedback on the manuscript. C.N., N.B., H.K., J.-Y.K., D.B.K., N.S., Z.N., R.L., J.H.K., G.C., H.L., and B.h.L. reviewed the manuscript and provided additional comments and/or editorial suggestions.

## DATA AND MATERIALS AVAILABILITY

MS-based protein identifications are available through the Plant Proteome Data Base (PPDB) under experiment ids #2599-2604. MS-based protein quantifications are available in the Supplemental Data Sets. RNA-seq data are available in the GEO repository under accession number GSE307303

## SUPPLEMENTAL INFORMATION

Supplemental Figures 1-5

Supplemental Tables 1-3

Supplemental Data Sets 1 and 2

## Supplemental Figures

**Supplemental Figure 1.**
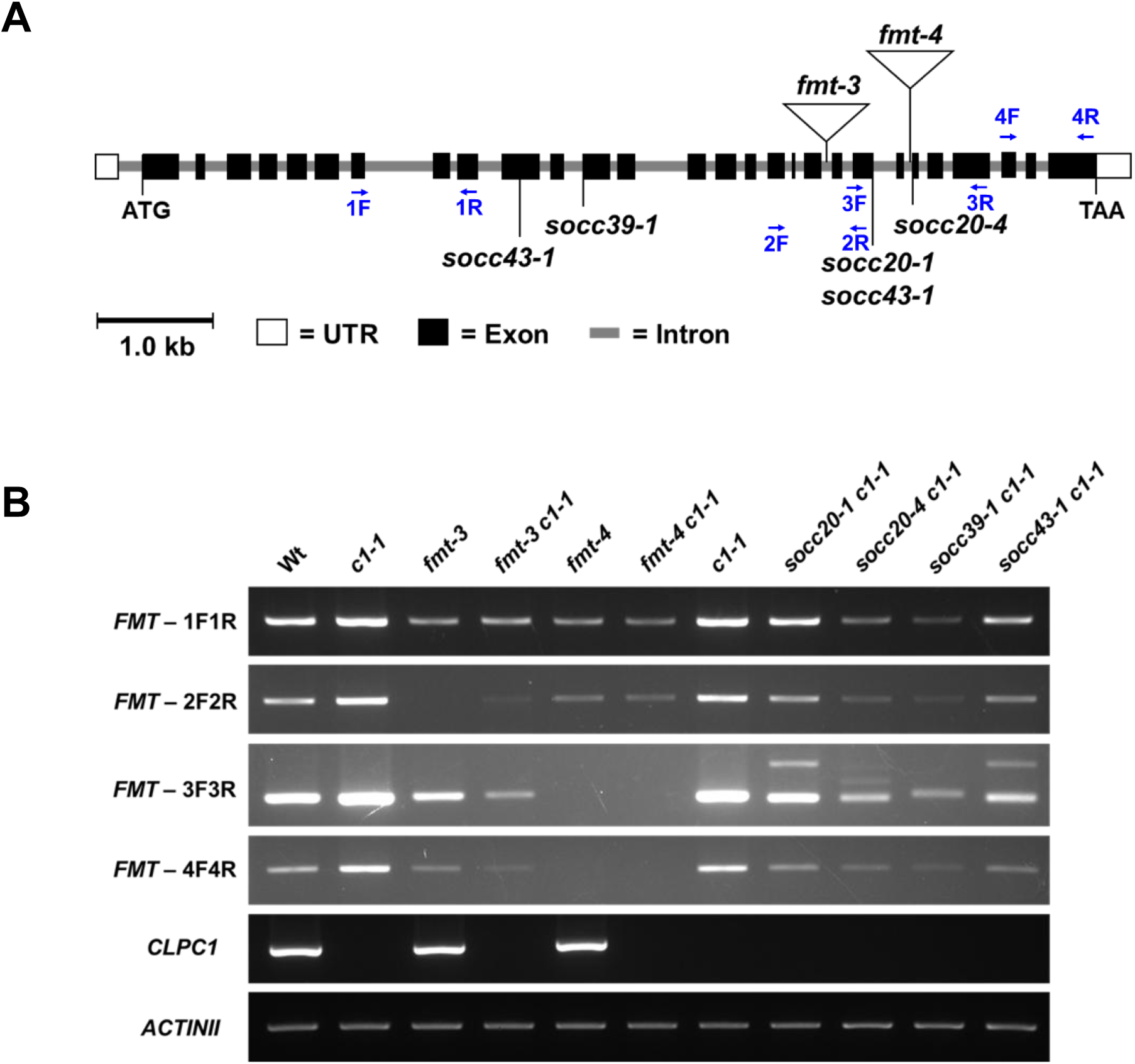
Analysis of *FMT* and *CLPC1* transcript accumulation by semi-quantitative RT-PCR. **(A)** *FMT* (At3g52140) gene model illustrating the positions of RT-PCR primer sets used in this study: *FMT*-1F1R (exons 7–9), *FMT*-2F2R (exons 17–21), *FMT*-3F3R (exons 21–25), and *FMT*-4F4R (exons 26–28). **(B)** Semi-quantitative RT-PCR analysis of *FMT* and *CLPC1* transcript levels in Col-0, *clpc1-1*, *fmt-3*, *fmt-3 clpc1-1*, *fmt-4*, *fmt-4 clpc1-1*, *socc20-1 clpc1-1*, *socc20-4 clpc1-1*, *socc39-1 clpc1-1*, and *socc43-1 clpc1-1*. *ACTINII* was used as a control. *ACTINII* was amplified for 22 cycles, while all other targets were amplified for 25 cycles. *FMT* expression was reduced in all *fmt* alleles, with complete loss in *fmt-4*, while *CLPC1* transcripts were absent in all *clpc1-1* backgrounds.

**Supplemental Figure 2.**
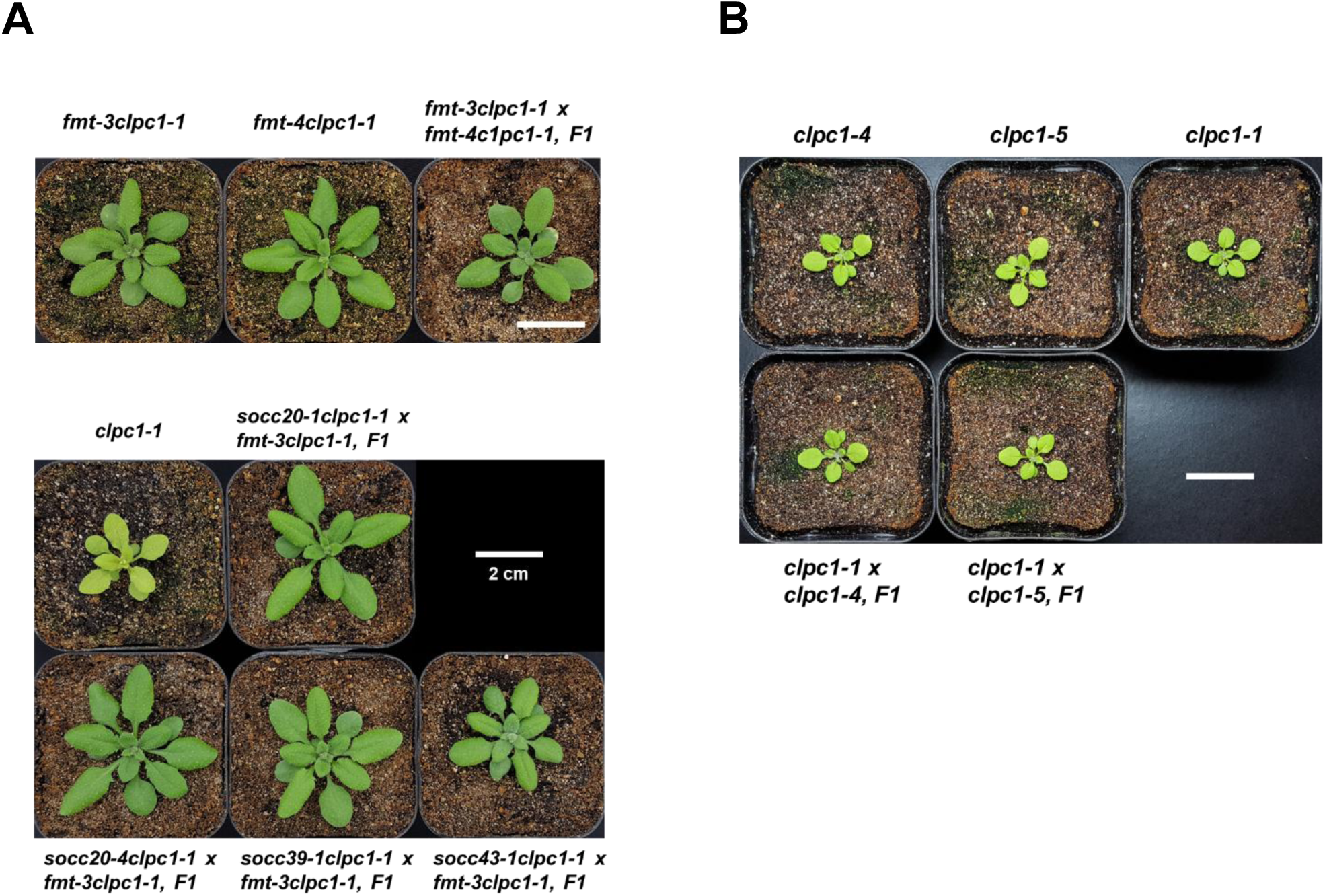
Heteroallelic analysis confirms that EMS-derived suppressors and CRISPR alleles affect *FMT* and *CLPC1*, respectively. **(A)** Heteroallelic crosses demonstrate that EMS-derived suppressors are allelic to *FMT*. Top panel: Growth and development of *fmt-3 clpc1-1*, *fmt-4 clpc1-1*, and F1 progeny from the *fmt-3 clpc1-1 × fmt-4 clpc1-1* cross. Plants were grown on soil for 22 days under long-day conditions (16/8 hour light/dark cycle at 100 µmol photons m⁻² s⁻¹). Scale bar = 2 cm. Bottom panel: Growth and development of *clpc1-1* and F1 progeny from heteroallelic crosses between *fmt-3 clpc1-1* and each EMS-derived suppressor: *socc20-1 clpc1-1*, *socc20-4 clpc1-1*, *socc39-1 clpc1-1*, and *socc43-1 clpc1-1*. Plants were grown under identical conditions. Scale bar = 2 cm. **(B)** CRISPR/Cas9-derived *clpc1* alleles phenocopy *clpc1-1* in homozygous and heteroallelic combinations. Growth and development of *clpc1-1*, *clpc1-4*, *clpc1-5*, and heteroallelic F1 plants from *clpc1-1 × clpc1-4* and *clpc1-1 × clpc1-5* crosses. All plants were grown on soil for 16 days under long-day conditions (16/8 hour light/dark cycle at 100 µmol photons m⁻² s⁻¹). Scale bar = 2 cm.

**Supplemental Figure 3.**
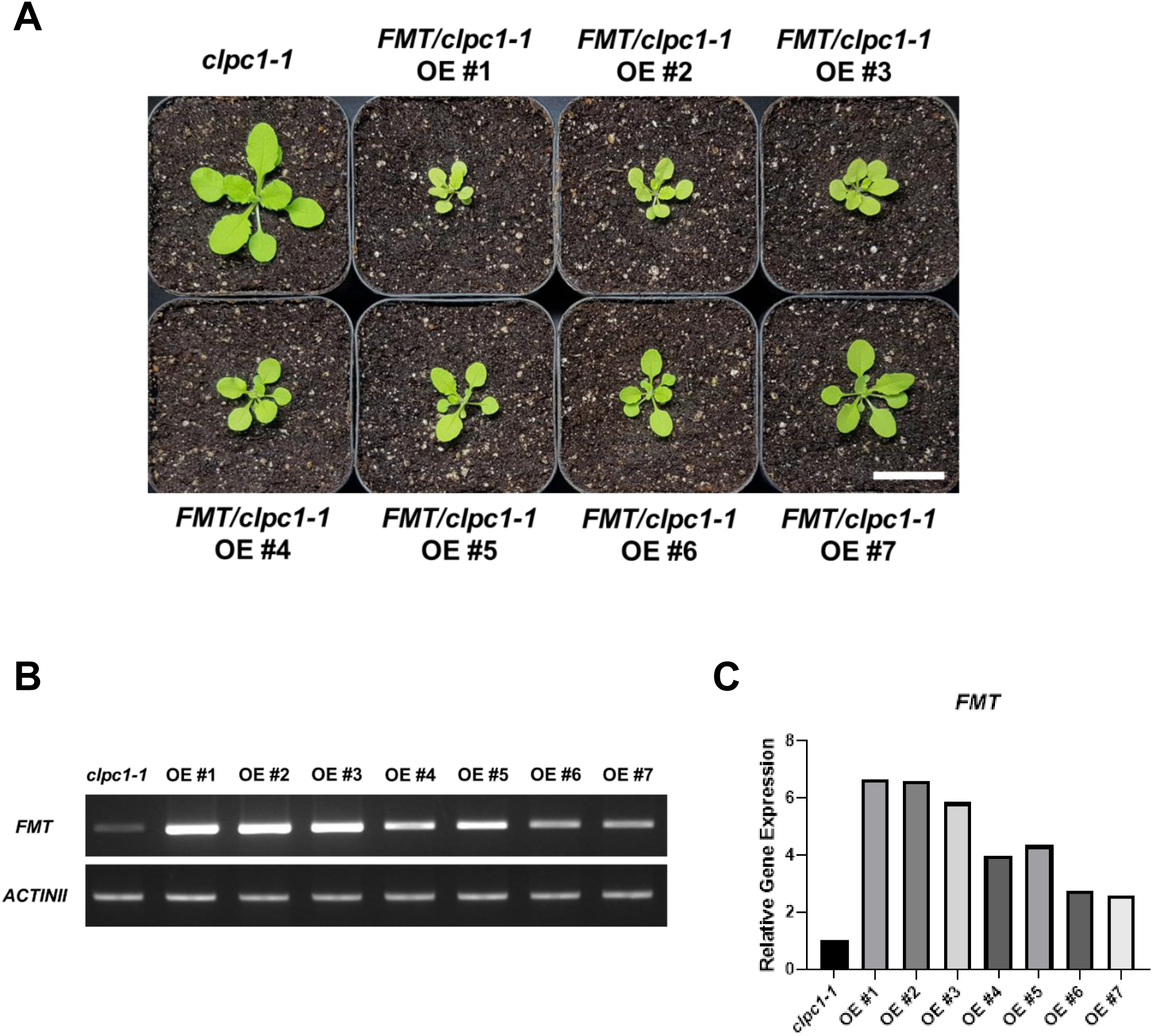
Overexpression of *FMT* in *clpc1-1* results in smaller, chlorotic plants that scale with transcript abundance. **(A)** Growth phenotype of *clpc1-1* and independent *FMT/clpc1-1* overexpression (OE) lines (#1–7) grown on soil for 21 days under a 16/8 hour light/dark cycle at 100 µmol photons m⁻² s⁻¹. Lines with stronger *FMT* expression exhibit smaller rosettes and more pronounced chlorosis (e.g., OE #1 shows stronger yellowing compared to OE #7). **(B)** Semi-quantitative RT-PCR analysis of *FMT* transcript levels in the same lines shown in (A). Primers targeting *FMT-RT3* (exons 21–25) were used to detect *FMT* transcript abundance. *ACTINII* served as loading control and was amplified for 22 cycles, whereas *FMT* was amplified for 25 cycles. **(C)** Quantification of *FMT* transcript abundance from (B). Both reduced plant size and chlorosis correlate negatively with *FMT* transcript levels across independent OE lines.

**Supplemental Figure 4.**
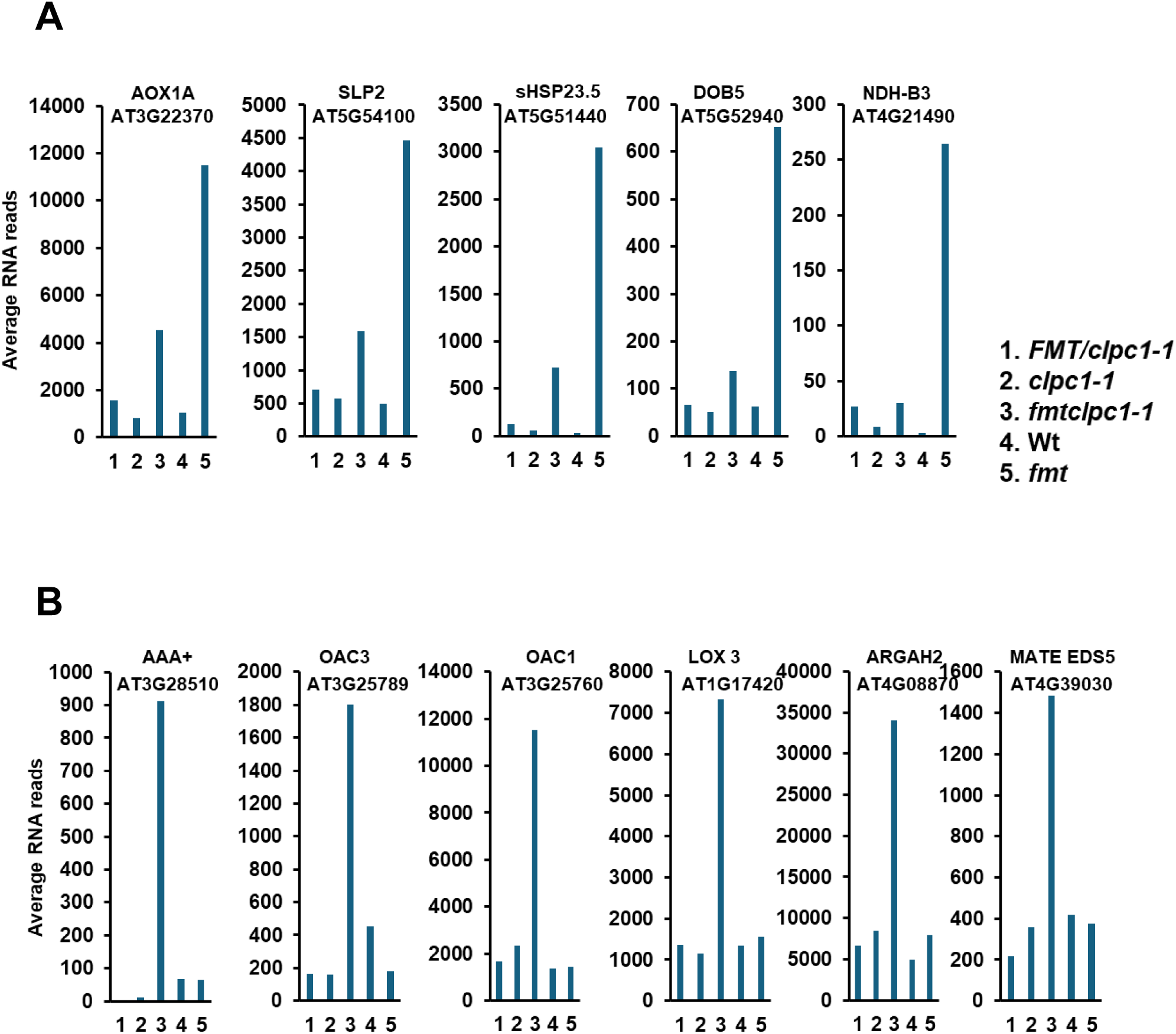
RNA expression patterns of selected genes in subclusters 4, 8, and 9 across five genotypes. RNA expression levels are shown for *clpc1-1*, *FMT/clpc1-1*, *fmtclpc1-1*, *fmt*, and WT. **(A, B)** Representative genes from subcluster 4 (within cluster II) and subclusters 8 and 9 (within cluster III) exhibiting high mRNA abundance in *fmt* (subcluster 4) or *fmtclpc1* (subclusters 8 and 9) relative to the other four genotypes. **(A)** Five genes in subcluster 4 for which RNA levels in *fmt* were >3-fold higher than in each of the other genotypes. **(B)** Six genes in subclusters 8 and 9 for which RNA levels in *fmtclpc1* were >3-fold higher than in each of the other genotypes.

**Supplemental Figure 5.**
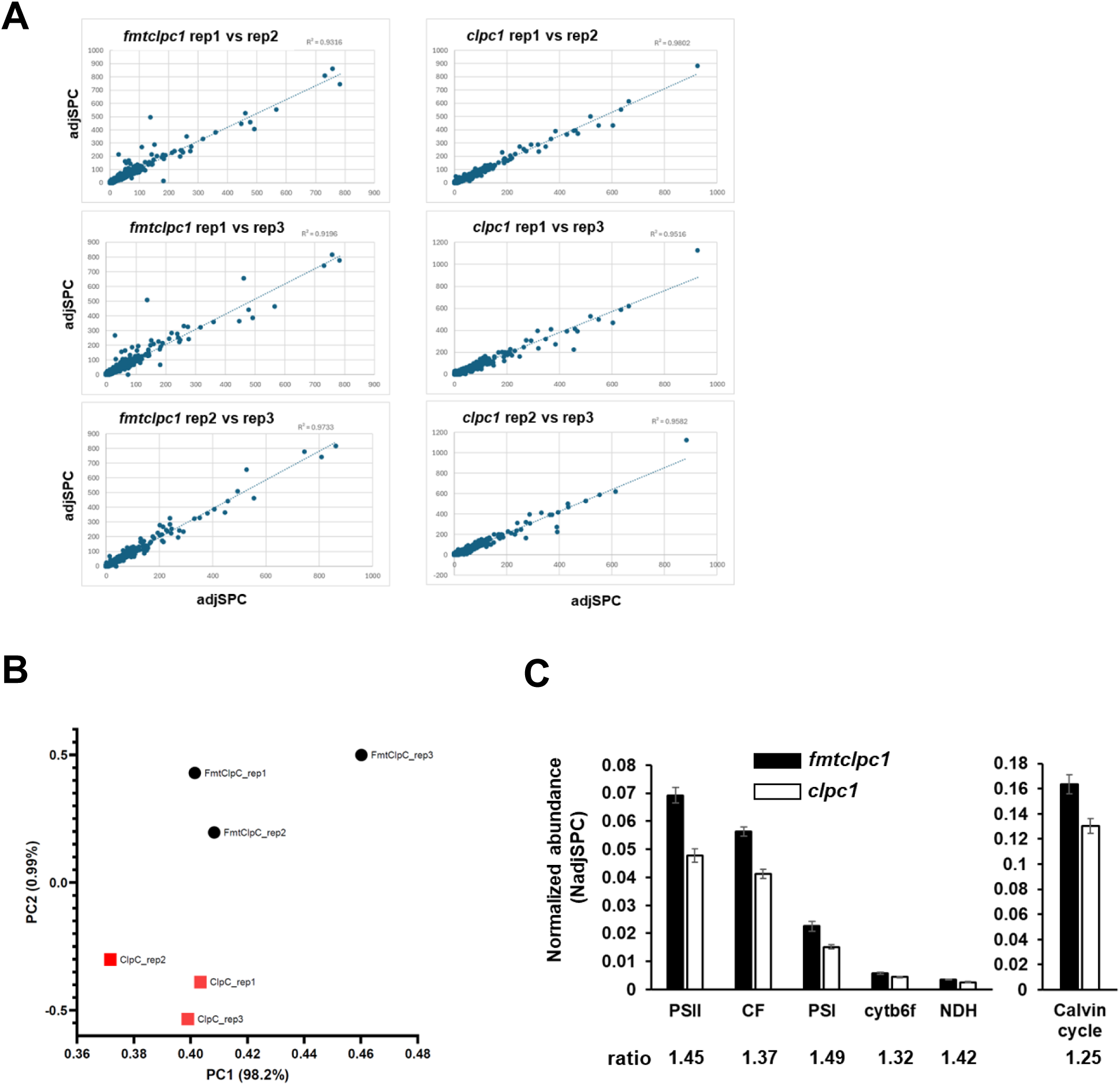
Reproducibility of biological proteomics replicates and protein investments in the photosynthetic apparatus. (**A**) Cross-correlation between biological replicates for each genotype based on AdjSPC (no normalization). For clarity, the X- and Y-axis are truncated at 1000, thus removing the very high data points for RBCL (between 2176 and 3105). (**B**) Principal component analysis (PCA) of the 3 biological samples for each genotype, illustrating reproducibility between replicates and separation of the two genotypes (*fmtclpc1* and *clpc1*). **(C)** Relative protein investments in the five complexes of the photosynthetic electron transport chain (Photosystem II (PSII), Photosystem I (PSI), the cytochrome b6f complex (cytb6f), NDH complex (NDH) and ATP synthase (CF), as well as proteins of the Calvin-Benson cycle. The *fmtclpc1/clpc1* ratios are indicated for each complex.

---

**Supplemental Table 1.**
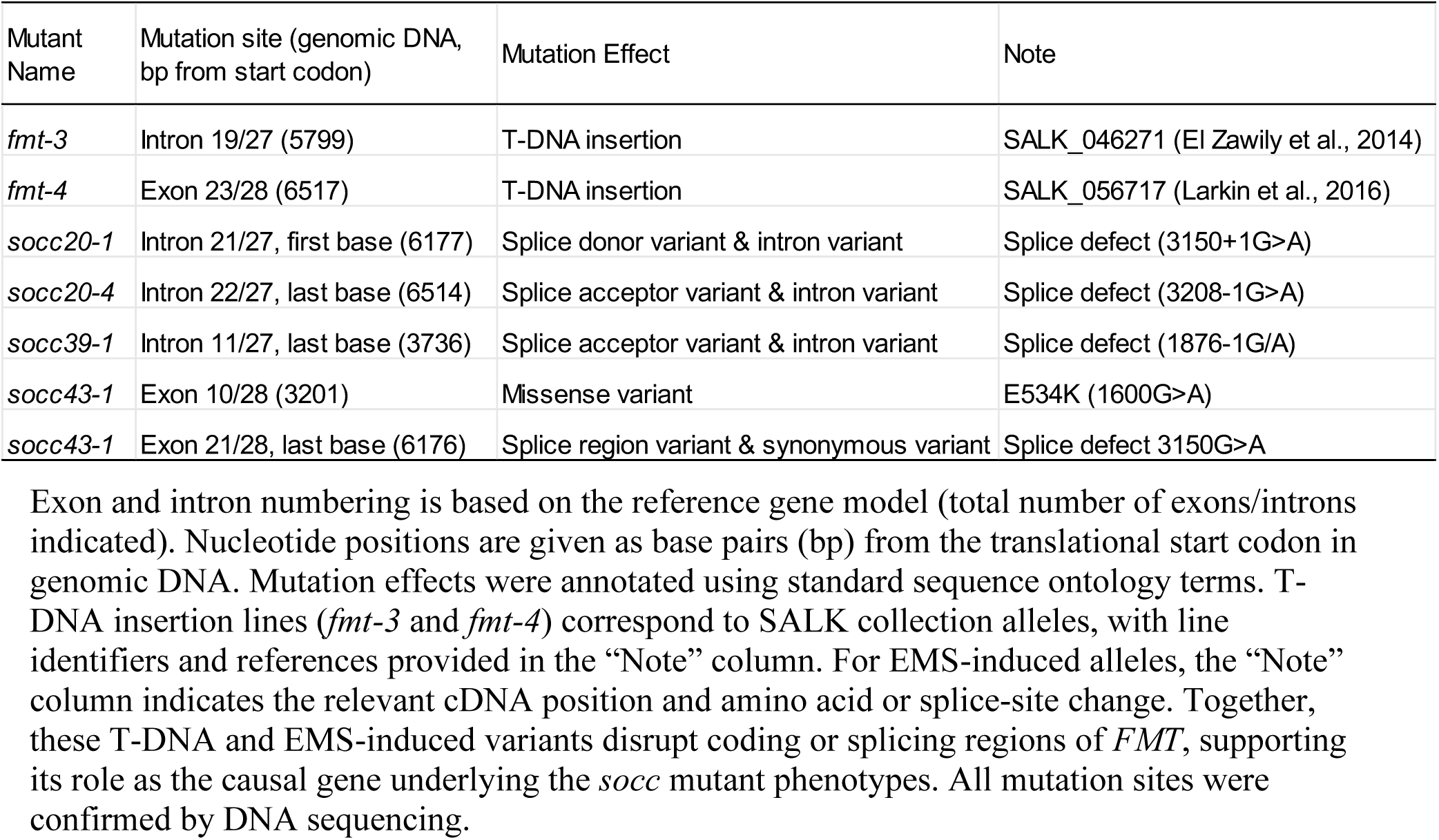
FMT lines generated and/or used in this study.

**Supplemental Table 2.**
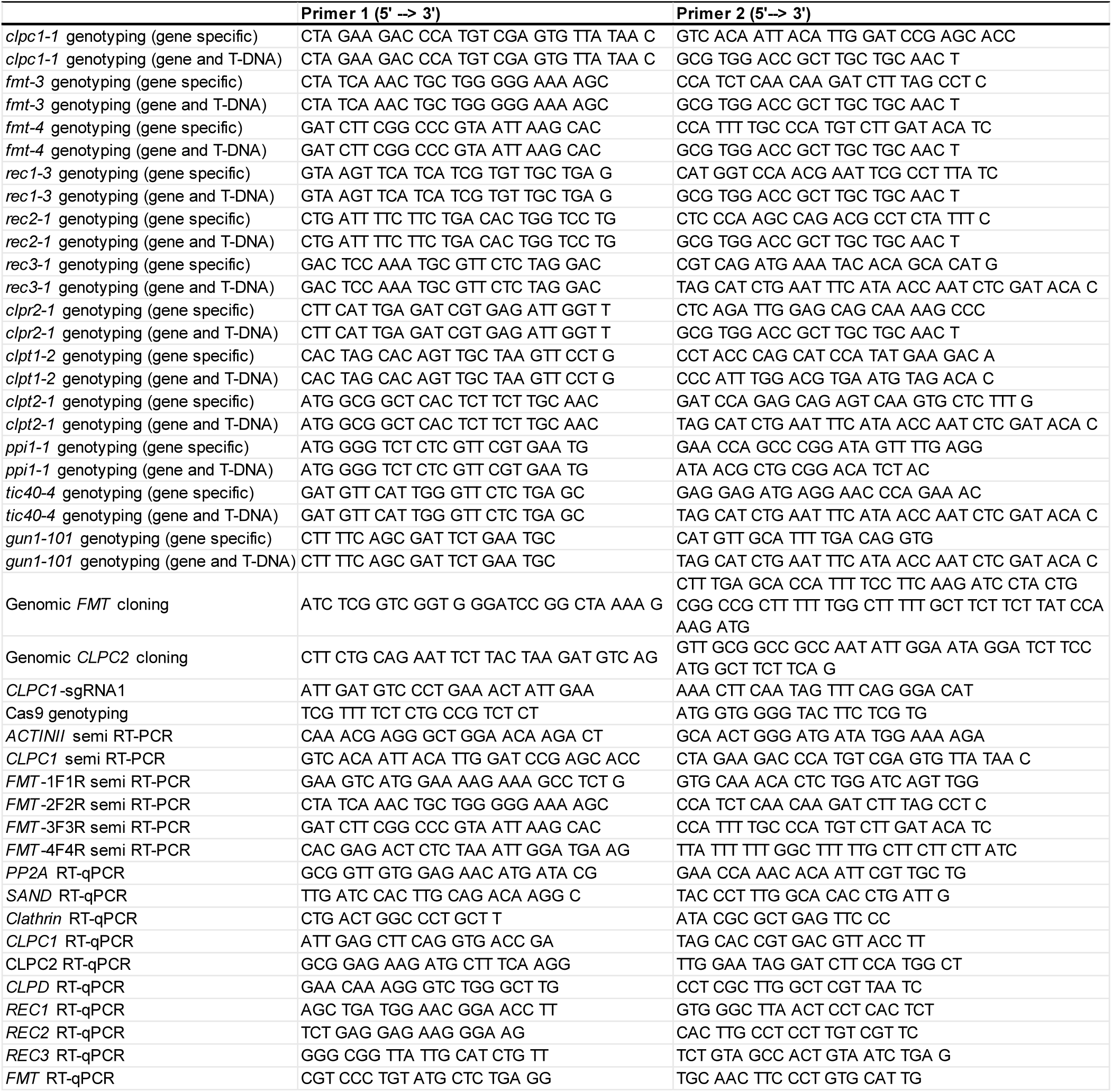
Primers used in this paper.

## REFERENCES

Atanasov, V., Schumacher, J., Muino, J.M., Larasati, C., Wang, L., Kaufmann, K., Leister, D., and Kleine, T. (2024). Arabidopsis BBX14 is involved in high light acclimation and seedling development. Plant J 118:141–158. 10.1111/tpj.16597.

Ayabe, H., Kawai, N., Shibamura, M., Fukao, Y., Fujimoto, M., Tsutsumi, N., and Arimura, S.I. (2021). FMT, a protein that affects mitochondrial distribution, interacts with translation-related proteins in Arabidopsis thaliana. Plant Cell Rep 40:327–337. 10.1007/s00299-020-02634-9.

Bae, N., Shim, S.H., Alavilli, H., Do, H., Park, M., Lee, D.W., Lee, J.H., Lee, H., Li, X., Lee, C.H., et al. (2025). Enhanced salt stress tolerance in plants without growth penalty through increased photosynthesis activity by plastocyanin from Antarctic moss. Plant J 121:e17168. 10.1111/tpj.17168.

Bedard, J., Trosch, R., Wu, F., Ling, Q., Flores-Perez, U., Topel, M., Nawaz, F., and Jarvis, P. (2017). Suppressors of the Chloroplast Protein Import Mutant tic40 Reveal a Genetic Link between Protein Import and Thylakoid Biogenesis. Plant Cell 29:1726–1747. 10.1105/tpc.16.00962.

Bittner, A., Ciesla, A., Gruden, K., Lukan, T., Mahmud, S., Teige, M., Vothknecht, U.C., and Wurzinger, B. (2022). Organelles and phytohormones: a network of interactions in plant stress responses. J Exp Bot 73:7165–7181. 10.1093/jxb/erac384.

Bouchnak, I., and van Wijk, K.J. (2021). Structure, function, and substrates of Clp AAA+ protease systems in cyanobacteria, plastids, and apicoplasts: A comparative analysis. J Biol Chem 296:100338. 10.1016/j.jbc.2021.100338.

Brownfield, D.L., Todd, C.D., and Deyholos, M.K. (2008). Analysis of Arabidopsis arginase gene transcription patterns indicates specific biological functions for recently diverged paralogs. Plant Mol Biol 67:429–440. 10.1007/s11103-008-9336-2.

Chen, Y., Chen, Y., Shi, C., Huang, Z., Zhang, Y., Li, S., Li, Y., Ye, J., Yu, C., Li, Z., et al. (2018). SOAPnuke: a MapReduce acceleration-supported software for integrated quality control and preprocessing of high-throughput sequencing data. Gigascience 7:1–6. 10.1093/gigascience/gix120.

Chustecki, J.M., Etherington, R.D., Gibbs, D.J., and Johnston, I.G. (2022). Altered collective mitochondrial dynamics in the Arabidopsis msh1 mutant compromising organelle DNA maintenance. J Exp Bot 73:5428–5439. 10.1093/jxb/erac250.

El Zawily, A.M., Schwarzlander, M., Finkemeier, I., Johnston, I.G., Benamar, A., Cao, Y., Gissot, C., Meyer, A.J., Wilson, K., Datla, R., et al. (2014). FRIENDLY regulates mitochondrial distribution, fusion, and quality control in Arabidopsis. Plant Physiol 166:808–828. 10.1104/pp.114.243824.

Elhafez, D., Murcha, M.W., Clifton, R., Soole, K.L., Day, D.A., and Whelan, J. (2006). Characterization of mitochondrial alternative NAD(P)H dehydrogenases in Arabidopsis: intraorganelle location and expression. Plant Cell Physiol 47:43–54. 10.1093/pcp/pci221.

Friso, G., Olinares, P.D., and van Wijk, K.J. (2011). The workflow for quantitative proteome analysis of chloroplast development and differentiation, chloroplast mutants, and protein interactions by spectral counting. Methods Mol Biol 775:265–282. 10.1007/978-1-61779-237-3_14.

Furumoto, T., Yamaguchi, T., Ohshima-Ichie, Y., Nakamura, M., Tsuchida-Iwata, Y., Shimamura, M., Ohnishi, J., Hata, S., Gowik, U., Westhoff, P., et al. (2011). A plastidial sodium-dependent pyruvate transporter. Nature 476:472–475. 10.1038/nature10250.

Gao, L.L., Hong, Z.H., Wang, Y., and Wu, G.Z. (2023). Chloroplast proteostasis: A story of birth, life, and death. Plant Commun 4:100424. 10.1016/j.xplc.2022.100424.

Gao, Y., and Zhao, Y. (2013). Epigenetic suppression of T-DNA insertion mutants in Arabidopsis. Mol Plant 6:539–545. 10.1093/mp/sss093.

Gehl, B., and Sweetlove, L.J. (2014). Mitochondrial Band-7 family proteins: scaffolds for respiratory chain assembly? Front Plant Sci 5:141. 10.3389/fpls.2014.00141.

Gehl, B., Lee, C.P., Bota, P., Blatt, M.R., and Sweetlove, L.J. (2014). An Arabidopsis stomatin-like protein affects mitochondrial respiratory supercomplex organization. Plant Physiol 164:1389–1400. 10.1104/pp.113.230383.

He, C., Berkowitz, O., Hu, S., Zhao, Y., Qian, K., Shou, H., Whelan, J., and Wang, Y. (2023). Co-regulation of mitochondrial and chloroplast function: Molecular components and mechanisms. Plant Commun 4:100496. 10.1016/j.xplc.2022.100496.

Hemono, M., Salinas-Giege, T., Roignant, J., Vingadassalon, A., Hammann, P., Ubrig, E., Ngondo, P., and Duchene, A.M. (2022). FRIENDLY (FMT) is an RNA binding protein associated with cytosolic ribosomes at the mitochondrial surface. Plant J 112:309–321. 10.1111/tpj.15962.

Hernandez-Verdeja, T. (2025). Regulation of Chloroplast Biogenesis and Differentiation. J Exp Bot 10.1093/jxb/eraf530.

Hu, Q., Zhang, H., Song, Y., Song, L., Zhu, L., Kuang, H., and Larkin, R.M. (2024). REDUCED CHLOROPLAST COVERAGE proteins are required for plastid proliferation and carotenoid accumulation in tomato. Plant Physiol 196:511–534. 10.1093/plphys/kiae275.

Jia, X., Chanda, B., Zhao, M., Brunner, A.M., and Beers, E.P. (2015). Instability of the Arabidopsis mutant csn5a-2 caused by epigenetic modification of intronic T-DNA. Plant Sci 238:53–63. 10.1016/j.plantsci.2015.05.015.

Kacprzak, S.M., and Van Aken, O. (2023). FRIENDLY is required for efficient dark-induced mitophagy and controlled senescence in Arabidopsis. Free Radic Biol Med 204:1–7. 10.1016/j.freeradbiomed.2023.04.007.

Kim, J., Rudella, A., Ramirez Rodriguez, V., Zybailov, B., Olinares, P.D., and van Wijk, K.J. (2009). Subunits of the plastid ClpPR protease complex have differential contributions to embryogenesis, plastid biogenesis, and plant development in Arabidopsis. Plant Cell 21:1669–1692. 10.1105/tpc.108.063784.

Kim, J., Olinares, P.D., Oh, S.H., Ghisaura, S., Poliakov, A., Ponnala, L., and van Wijk, K.J. (2013). Modified Clp protease complex in the ClpP3 null mutant and consequences for chloroplast development and function in Arabidopsis. Plant Physiol 162:157–179. 10.1104/pp.113.215699.

Kim, J., Kimber, M.S., Nishimura, K., Friso, G., Schultz, L., Ponnala, L., and van Wijk, K.J. (2015). Structures, Functions, and Interactions of ClpT1 and ClpT2 in the Clp Protease System of Arabidopsis Chloroplasts. Plant Cell 27:1477–1496. 10.1105/tpc.15.00106.

Kim, Y., Schumaker, K.S., and Zhu, J.K. (2006). EMS mutagenesis of Arabidopsis. Methods Mol Biol 323:101–103. 10.1385/1-59745-003-0:101.

Kovacheva, S., Bédard, J., Wardle, A., Patel, R., and Jarvis, P. (2007). Further in vivo studies on the role of the molecular chaperone, Hsp93, in plastid protein import. Plant J. 50:364–379. 10.1111/j.1365-313X.2007.03060.x.

Kubis, S., Baldwin, A., Patel, R., Razzaq, A., Dupree, P., Lilley, K., Kurth, J., Leister, D., and Jarvis, P. (2003). The Arabidopsis ppi1 mutant is specifically defective in the expression, chloroplast import, and accumulation of photosynthetic proteins. Plant Cell 15:1859–1871. 10.1105/tpc.012955.

Lama, S., Broda, M., Abbas, Z., Vaneechoutte, D., Belt, K., Sall, T., Vandepoele, K., and Van Aken, O. (2019). Neofunctionalization of Mitochondrial Proteins and Incorporation into Signaling Networks in Plants. Mol Biol Evol 36:974–989. 10.1093/molbev/msz031.

Larkin, R.M., Stefano, G., Ruckle, M.E., Stavoe, A.K., Sinkler, C.A., Brandizzi, F., Malmstrom, C.M., and Osteryoung, K.W. (2016). REDUCED CHLOROPLAST COVERAGE genes from Arabidopsis thaliana help to establish the size of the chloroplast compartment. Proc Natl Acad Sci U S A 113:E1116–1125. 10.1073/pnas.1515741113.

Lee, K.P., Li, M., Li, M., Liu, K., Medina-Puche, L., Qi, S., Cui, C., Lozano-Duran, R., and Kim, C. (2023). Hierarchical regulatory module GENOMES UNCOUPLED1-GOLDEN2-LIKE1/2-WRKY18/40 modulates salicylic acid signaling. Plant Physiol 192:3120–3133. 10.1093/plphys/kiad251.

Liao, Y., Smyth, G.K., and Shi, W. (2014). featureCounts: an efficient general purpose program for assigning sequence reads to genomic features. Bioinformatics 30:923–930. 10.1093/bioinformatics/btt656.

Llamas, E., and Pulido, P. (2022). A proteostasis network safeguards the chloroplast proteome. Essays Biochem 66:219–228. 10.1042/EBC20210058.

Logan, D.C., Scott, I., and Tobin, A.K. (2003). The genetic control of plant mitochondrial morphology and dynamics. Plant J 36:500–509. 10.1046/j.1365-313x.2003.01894.x.

Lopez, B., Izquierdo, Y., Cascon, T., Zamarreno, A.M., Garcia-Mina, J.M., Pulido, P., and Castresana, C. (2024). Mutant noxy8 exposes functional specificities between the chloroplast chaperones CLPC1 and CLPC2 in the response to organelle stress and plant defence. Plant Cell Environ 47:2336–2350. 10.1111/pce.14882.

Love, M.I., Huber, W., and Anders, S. (2014). Moderated estimation of fold change and dispersion for RNA-seq data with DESeq2. Genome Biol 15:550. 10.1186/s13059-014-0550-8.

Ma, J., Liang, Z., Zhao, J., Wang, P., Ma, W., Mai, K.K., Fernandez Andrade, J.A., Zeng, Y., Grujic, N., Jiang, L., et al. (2021). Friendly mediates membrane depolarization-induced mitophagy in Arabidopsis. Curr Biol 31:1931–1944 e1934. 10.1016/j.cub.2021.02.034.

Marty, L., Bausewein, D., Muller, C., Bangash, S.A.K., Moseler, A., Schwarzlander, M., Muller-Schussele, S.J., Zechmann, B., Riondet, C., Balk, J., et al. (2019). Arabidopsis glutathione reductase 2 is indispensable in plastids, while mitochondrial glutathione is safeguarded by additional reduction and transport systems. New Phytol 224:1569–1584. 10.1111/nph.16086.

Morikawa, K., Shiina, T., Murakami, S., and Toyoshima, Y. (2002). Novel nuclear-encoded proteins interacting with a plastid sigma factor, Sig1, in Arabidopsis thaliana. FEBS Lett 514:300–304. 10.1016/s0014-5793(02)02388-8.

Nakamura, S., Hagihara, S., Otomo, K., Ishida, H., Hidema, J., Nemoto, T., and Izumi, M. (2021). Autophagy Contributes to the Quality Control of Leaf Mitochondria. Plant Cell Physiol 62:229–247. 10.1093/pcp/pcaa162.

Nishimura, K., Apitz, J., Friso, G., Kim, J., Ponnala, L., Grimm, B., and van Wijk, K.J. (2015). Discovery of a Unique Clp Component, ClpF, in Chloroplasts: A Proposed Binary ClpF-ClpS1 Adaptor Complex Functions in Substrate Recognition and Delivery. Plant Cell 27:2677–2691. 10.1105/tpc.15.00574.

Nishimura, K., Asakura, Y., Friso, G., Kim, J., Oh, S.H., Rutschow, H., Ponnala, L., and van Wijk, K.J. (2013). ClpS1 is a conserved substrate selector for the chloroplast Clp protease system in Arabidopsis. Plant Cell 25:2276–2301. 10.1105/tpc.113.112557.

Ohmiya, A., Oda-Yamamizo, C., and Kishimoto, S. (2019). Overexpression of CONSTANS-like 16 enhances chlorophyll accumulation in petunia corollas. Plant Sci 280:90–96. 10.1016/j.plantsci.2018.11.013.

Olinares, P.D., Kim, J., Davis, J.I., and van Wijk, K.J. (2011). Subunit stoichiometry, evolution, and functional implications of an asymmetric plant plastid ClpP/R protease complex in Arabidopsis. Plant Cell 23:2348–2361. 10.1105/tpc.111.086454.

Paila, Y.D., Richardson, L.G.L., and Schnell, D.J. (2015). New insights into the mechanism of chloroplast protein import and its integration with protein quality control, organelle biogenesis and development. J Mol Biol 427:1038–1060. 10.1016/j.jmb.2014.08.016.

Rei Liao, J.Y., Friso, G., Forsythe, E.S., Michel, E.J.S., Williams, A.M., Boguraev, S.S., Ponnala, L., Sloan, D.B., and van Wijk, K.J. (2022). Proteomics, phylogenetics, and coexpression analyses indicate novel interactions in the plastid CLP chaperone-protease system. J Biol Chem 298:101609. 10.1016/j.jbc.2022.101609.

Rekhter, D., Lüdke, D., Ding, Y., Feussner, K., Zienkiewicz, K., Lipka, V., Wiermer, M., Zhang, Y., and Feussner, I. (2019). Isochorismate-derived biosynthesis of the plant stress hormone salicylic acid. Science 365:498–502. 10.1126/science.aaw1720.

Richter, A.S., Nagele, T., Grimm, B., Kaufmann, K., Schroda, M., Leister, D., and Kleine, T. (2023). Retrograde signaling in plants: A critical review focusing on the GUN pathway and beyond. Plant Commun 4:100511. 10.1016/j.xplc.2022.100511.

Rodriguez-Concepcion, M., D’Andrea, L., and Pulido, P. (2019). Control of plastidial metabolism by the Clp protease complex. J Exp Bot 70:2049–2058. 10.1093/jxb/ery441.

Rudella, A., Friso, G., Alonso, J.M., Ecker, J.R., and van Wijk, K.J. (2006). Downregulation of ClpR2 leads to reduced accumulation of the ClpPRS protease complex and defects in chloroplast biogenesis in Arabidopsis. Plant Cell 18:1704–1721. 10.1105/tpc.106.042861.

Selinski, J., Scheibe, R., Day, D.A., and Whelan, J. (2018). Alternative Oxidase Is Positive for Plant Performance. Trends Plant Sci 23:588–597. 10.1016/j.tplants.2018.03.012.

Sjogren, L.L., and Clarke, A.K. (2011). Assembly of the chloroplast ATP-dependent Clp protease in Arabidopsis is regulated by the ClpT accessory proteins. Plant Cell 23:322–332. 10.1105/tpc.110.082321.

Sjogren, L.L., Stanne, T.M., Zheng, B., Sutinen, S., and Clarke, A.K. (2006). Structural and functional insights into the chloroplast ATP-dependent Clp protease in Arabidopsis. Plant Cell 18:2635–2649. 10.1105/tpc.106.044594.

Sun, J.L., Li, J.Y., Wang, M.J., Song, Z.T., and Liu, J.X. (2021). Protein Quality Control in Plant Organelles: Current Progress and Future Perspectives. Mol Plant 14:95–114. 10.1016/j.molp.2020.10.011.

Sun, Y., and Jarvis, R.P. (2023). Chloroplast Proteostasis: Import, Sorting, Ubiquitination, and Proteolysis. Annu Rev Plant Biol 74:259–283. 10.1146/annurev-arplant-070122-032532.

Susila, H., Nasim, Z., Gawarecka, K., Jung, J.Y., Jin, S., Youn, G., and Ahn, J.H. (2023). Chloroplasts prevent precocious flowering through a GOLDEN2-LIKE-B-BOX DOMAIN PROTEIN module. Plant Commun 4:100515. 10.1016/j.xplc.2023.100515.

Tang, Q., Xu, D., Lenzen, B., Brachmann, A., Yapa, M.M., Doroodian, P., Schmitz-Linneweber, C., Masuda, T., Hua, Z., Leister, D., et al. (2024). GENOMES UNCOUPLED PROTEIN1 binds to plastid RNAs and promotes their maturation. Plant Commun 5:101069. 10.1016/j.xplc.2024.101069.

Tapken, W., Kim, J., Nishimura, K., van Wijk, K.J., and Pilon, M. (2015). The Clp protease system is required for copper ion-dependent turnover of the PAA2/HMA8 copper transporter in chloroplasts. New Phytol 205:511–517. 10.1111/nph.13093.

Thimm, O., Blasing, O., Gibon, Y., Nagel, A., Meyer, S., Kruger, P., Selbig, J., Muller, L.A., Rhee, S.Y., and Stitt, M. (2004). MAPMAN: a user-driven tool to display genomics data sets onto diagrams of metabolic pathways and other biological processes. Plant J 37:914–939. 10.1111/j.1365-313x.2004.02016.x.

Tzvetkova-Chevolleau, T., Franck, F., Alawady, A.E., Dall’Osto, L., Carriere, F., Bassi, R., Grimm, B., Nussaume, L., and Havaux, M. (2007). The light stress-induced protein ELIP2 is a regulator of chlorophyll synthesis in Arabidopsis thaliana. Plant J 50:795–809. 10.1111/j.1365-313X.2007.03090.x.

van Wijk, K.J. (2024). Intra-chloroplast proteases: A holistic network view of chloroplast proteolysis. Plant Cell 36:3116–3130. 10.1093/plcell/koae178.

Wachsman, G., Modliszewski, J.L., Valdes, M., and Benfey, P.N. (2017). A SIMPLE Pipeline for Mapping Point Mutations. Plant Physiol 174:1307–1313. 10.1104/pp.17.00415.

Wang, Y., Berkowitz, O., Selinski, J., Xu, Y., Hartmann, A., and Whelan, J. (2018). Stress responsive mitochondrial proteins in Arabidopsis thaliana. Free Radic Biol Med 122:28–39. 10.1016/j.freeradbiomed.2018.03.031.

Wang, Y., Selinski, J., Mao, C., Zhu, Y., Berkowitz, O., and Whelan, J. (2020). Linking mitochondrial and chloroplast retrograde signalling in plants. Philos Trans R Soc Lond B Biol Sci 375:20190410. 10.1098/rstb.2019.0410.

Waters, E.R., and Vierling, E. (2020). Plant small heat shock proteins - evolutionary and functional diversity. New Phytol 227:24–37. 10.1111/nph.16536.

Woodson, J.D., and Chory, J. (2008). Coordination of gene expression between organellar and nuclear genomes. Nat Rev Genet 9:383–395. 10.1038/nrg2348.

Wu, G.Z., Meyer, E.H., Richter, A.S., Schuster, M., Ling, Q., Schottler, M.A., Walther, D., Zoschke, R., Grimm, B., Jarvis, R.P., et al. (2019). Control of retrograde signalling by protein import and cytosolic folding stress. Nat Plants 5:525–538. 10.1038/s41477-019-0415-y.

Xie, Y.D., Li, W., Guo, D., Dong, J., Zhang, Q., Fu, Y., Ren, D., Peng, M., and Xia, Y. (2010). The Arabidopsis gene SIGMA FACTOR-BINDING PROTEIN 1 plays a role in the salicylate- and jasmonate-mediated defence responses. Plant Cell Environ 33:828–839. 10.1111/j.1365-3040.2009.02109.x.

Zhang, H., Zhang, L., Ji, Y., Jing, Y., Li, L., Chen, Y., Wang, R., Zhang, H., Yu, D., and Chen, L. (2022). Arabidopsis SIGMA FACTOR BINDING PROTEIN1 (SIB1) and SIB2 inhibit WRKY75 function in abscisic acid-mediated leaf senescence and seed germination. J Exp Bot 73:182–196. 10.1093/jxb/erab391.

